# Identification of PIM1 substrates reveals a role for NDRG1 in prostate cancer cellular migration and invasion

**DOI:** 10.1101/2020.01.21.913962

**Authors:** Russell J. Ledet, Sophie Ruff, Yu Wang, Shruti Nayak, Jeffrey A. Schneider, Beatrix Ueberheide, Susan K. Logan, Michael J. Garabedian

## Abstract

PIM1 is an oncogenic serine/threonine kinase that promotes and maintains prostate tumorigenesis. To more fully understand the mechanism by which PIM1 promotes oncogenesis, we performed a chemical genetic screen to identify direct PIM1 substrates in prostate cancer cells. The PIM1 substrates we identified were involved in a variety of oncogenic processes, and included N-Myc Downstream-Regulated Gene 1 (NDRG1), which has reported roles in the suppression of cancer cell invasion and metastasis. NDRG1 is phosphorylated by PIM1 at serine 330 (pS330), and the level of NDRG1 pS330 is associated with high grade compared to low grade prostate tumors. While NDRG1 pS330 is largely cytoplasmic, total NDRG1 is both cytoplasmic and nuclear. Mechanistically, PIM1 phosphorylation of NDRG1 decreases its stability, reducing its interaction with AR, and thereby lowering expression of AR target genes. PIM1-dependent NDRG1 phosphorylation also reduces NDRG1’s ability to suppress prostate cancer cell migration and invasion. Our study identifies a novel set of PIM1 substrates in prostate cancer cells using a direct, unbiased chemical genetic screen. It also provides key insights into the mechanisms by which PIM1-mediated phosphorylation of NDRG1 impairs its function, resulting in enhanced cell migration and invasion.

## INTRODUCTION

Pro-viral integration site for Moloney murine leukemia virus-1 (PIM1) is a proto-oncogene encoding a serine/threonine kinase^1,2^. PIM1 is constitutively active and does not depend upon post-translational modifications for activation^3,4^. As such, PIM1 is regulated primarily by transcription, including induction by cytokines through the JAK/STAT pathway^5^ and by hypoxia^6^. PIM1 protein levels are also regulated translationally via its 5’ UTR^7^ and by the microRNA miR-33a^8^. PIM1 has been implicated as an oncogene in both hematological malignancies, such as large B cell lymphoma^9^, and cancers of epithelial origins, such as breast and prostate cancer^10–13^. Whereas absent or weak expression of PIM1 by immunohistochemistry are observed in most benign prostate samples, moderate to strong levels of PIM1 are evident in ∼ 50% of prostate cancers^12^.

In mouse models of prostate cancer, conditional overexpression of PIM1 in prostate epithelial cells results in prostate intraepithelial neoplasia^13^. PIM1 also cooperates with c-MYC, an established oncogene, to promote advanced prostate adenocarcinoma^10^. We and others have previously shown that PIM1 phosphorylates the androgen receptor (AR)^14^, the main driver of prostate cancer and the main target in prostate cancer therapy^15,16^. PIM1-mediated AR serine 213 phosphorylation (pS213) differentially impacts AR target gene expression and is correlated with prostate cancer recurrence^14,17^. Given that deregulation of kinases is a hallmark of cancer^18^, we hypothesized that identifying PIM1 substrates and their phosphorylation sites in prostate cancer cells could help to elucidate PIM1’s role in the disease, and help to identify cancers with active PIM1.

To identify PIM1 substrates and their phosphorylation sites in LNCaP cells, we coupled a chemical genetic screen with a peptide capture, mass spectrometry (MS)-based approach^19^. We mutated the PIM1 gatekeeper residue in the ATP binding site to accept a bulky ATP analog. By using an ATP analog labeled with a thiol group on the γ-phosphate, we were able specifically label PIM1 substrates even in the presence of other cellular kinases ^20^.

A similar approach has revealed the direct substrates of AMPK, with unexpected roles in mitosis and cytokinesis^21,22^; CDK9, with functions in transcriptional termination through phosphorylation of the 5’-3’ exonuclease XRN2^23^; and CDK2, with a role in the DNA damage response by phosphorylation of NBS1, a necessary protein for DNA damage repair^24^. Overall, this is a rigorous approach that has revealed new substrates of kinases to yield novel mechanistic insights.

Several putative PIM1 substrates, including c-MYC, AR, and BAD, have been identified and found to be dysregulated in prostate cancer^14,17,25,26^. Still, the mechanism by which PIM1 promotes prostate cancer is not fully understood. Our study identifies new PIM1 substrates in prostate cancer cells in a direct, unbiased manner. This revealed that PIM1 substrates are involved in a variety of cellular processes, ranging from cell cycle checkpoints, to nucleic acid metabolism, to transcriptional regulation, to cellular motility and invasion. By further exploring the PIM1-dependent phosphorylation of NDRG1, we find that PIM1-dependent phosphorylation reduces the function of NDRG1 as a metastasis suppressor.

## RESULTS

### Identification of PIM1 substrates in prostate cancer cells

We conducted a chemical genetic screen to identify direct PIM1 substrates and their phosphorylation sites in LNCaP cells. We used an analog-sensitive (AS) kinase-substrate detection method^27,28^. To create the AS PIM1 kinase, we aligned the PIM1 amino acid sequence of the 33kDA PIM1 isoform^29^ with other kinases for which AS mutants have been generated, and identified leucine 120 (L120) as the conserved gatekeeper residue (Suppl. Table 1). We mutated L120 to a smaller glycine residue (L120G) and generated pools of LNCaP cells stably expressing similar levels of either wild type PIM1 (LNCaP-WT PIM1) or AS PIM1 (LNCaP-AS PIM1). We tested the ability of AS PIM1 to utilize a series of bulky N^6^-substituted ATP analogs in cells by treating with digitonin, a mild permeabilizing agent. We then lysed the cells and added *para*-nitrobenzyl mesylate (*p*NBM) to alkylate thiophosphorylated proteins, and used a thiophosphate ester-specific antibody to reveal substrates by Western blot (Fig. 1A). LNCaP cells harboring the AS PIM1 were able to utilize PhET-ATPγS, and to a lesser extent, Phe-ATPγS, to phosphorylate substrates. Cells expressing WT PIM1 were much less efficient at utilizing these analogs, whereas ATPγS is not specific for AS PIM1 (Fig. 1B). Therefore, all subsequent thiophosphorylation experiments were performed with PheET-ATPγS as the ATP analog.

**Figure 1.**
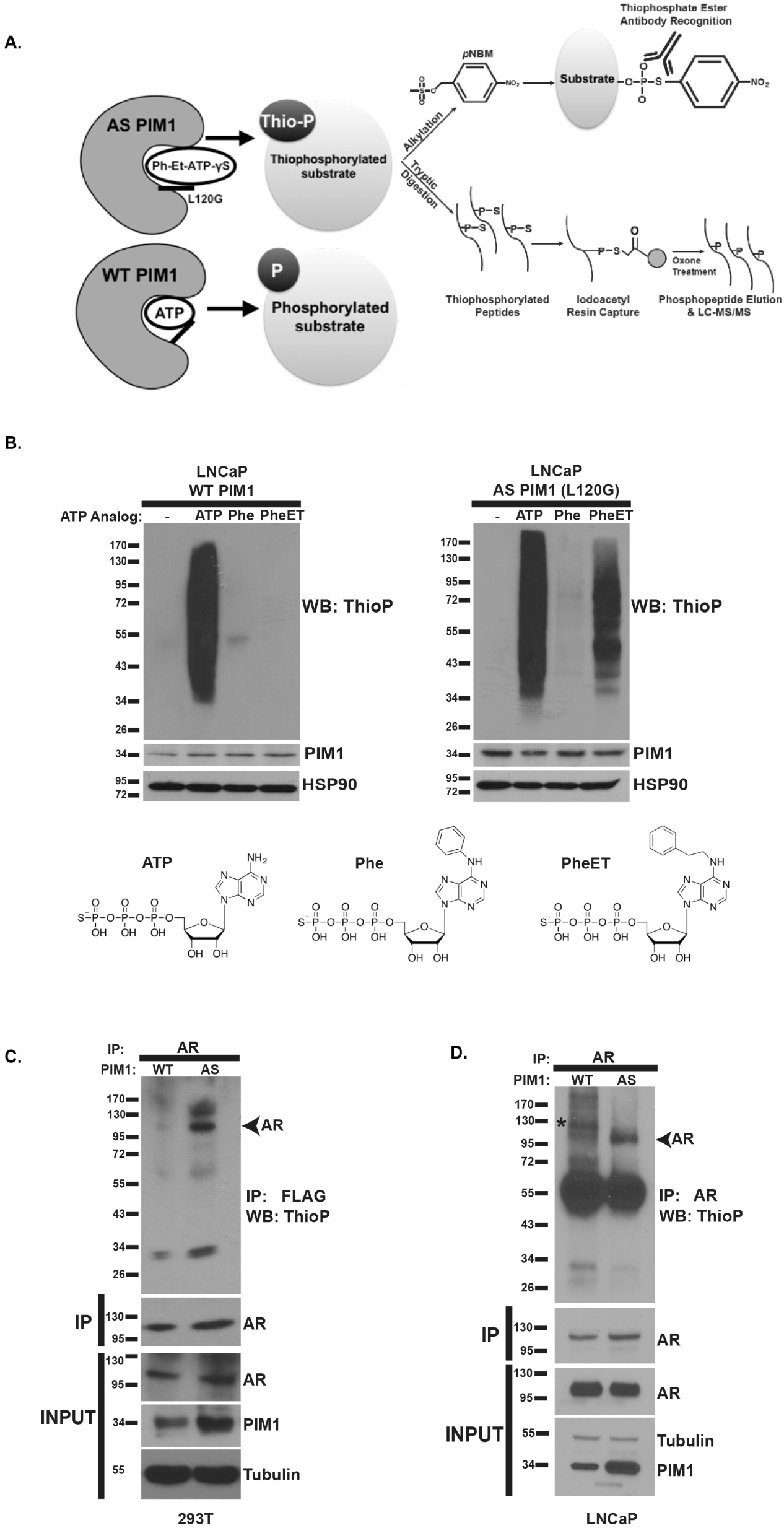
Screening Strategy to Identify PIM1 Substrates and Phosphorylation Sites in Prostate Cancer Cells. A) Schematic of the peptide-capture technique used to identify analog-sensitive (AS) PIM1 substrates and phosphorylation sites in prostate cancer cells. AS PIM1 uses ATPγS, a bulky ATP analog, to thiophosphorylate substrates. Upper panel: thiophosphorylated substrates are alkylated by p-nitrobenzyl mesylate (PNBM) and recognized by an antibody to the thiophosphate ester moiety (ThioP). Lower panel: thiophosphorylated peptides are captured on a resin, eluted, and identified using liquid chromatography-tandem mass spectrometry (LC-MS/MS). B) ATP analog optimization in LNCaP cells stably expressing either WT PIM1 or AS PIM1. Cells were treated with indicated ATP analog (ATP=ATPγS; Phe=N^6^-Phenyl-ATPγS; PheET=N^6^-Phenylethyl ATPγS) in the presence of digitonin, a mild-permeabilizing agent. Alkylation was completed using PNBM. Whole-cell lysates were analyzed by western blot for the presence of thiophosphorylation (ThioP), PIM1 (PIM1) and HSP90 as loading control. C) AS PIM1 thiophosphorylates androgen receptor (AR) expressed in 293T cells. FLAG-tagged WT AR was co-expressed with either WT PIM1 or AS PIM1 in 293T cells, treated with PhET ATPγS as above. AR was immunoprecipitated using FLAG (M2) agarose beads, and western blot was performed for thiophosphorylation (ThioP) and AR (FLAG). Input shows the protein levels of PIM1 and AR expressed in 293T cells, and immunoprecipitated AR. Tubulin serves as a loading control. D) AS PIM1 thiophosphorylates endogenous AR in LNCaP cells. LNCaP-WT PIM1 and LNCaP-AS PIM1 cells were treated PhET ATPγS as above. AR was immunoprecipitated and analyzed by western blot for the presence of thiophosphorylation (thioP) and AR. Asterisk denotes a non-specific background band. Input shows the protein levels in cells of PIM1, and endogenous AR, as well as the abundance of immunoprecipitated AR. Tubulin acts as a loading control. Western blots are representative of two independent experiments.

We have previously reported that AR is a substrate of PIM1^14^. To ensure that AS PIM1 retained its substrate specificity and could phosphorylate a known substrate, we immunoprecipitated ectopically expressed AR from 293T cells expressing either WT PIM1 or AS PIM1 and performed the thiophosphorylation assay. We also examined whether endogenous AR could be selectively thiophosphorylated in LNCaP-AS PIM1 as compared to LNCaP-WT PIM1 cells. Indeed, we found that AS PIM1 was able to use PhET-ATPγS to thiophosphorylate both ectopically expressed AR and endogenous AR in 293T and LNCaP cells, respectively. However, WT PIM1 was much less effective (Figs. 1C-D). This indicates that the AS PIM1 retains specificity for a known substrate.

To identify PIM1 substrates and their phosphorylation sites, we treated LNCaP-WT PIM1 and LNCaP-AS PIM1 expressing cells with PhET-ATPγS (Fig. 2A), and isolated thiophosphorylated tryptic-peptides using iodoacetyl beads^27^. This was followed by selective release of the thiophosphorylated peptides with spontaneous hydrolysis using potassium peroxomonosulfate (OXONE), forming phosphopeptides. Liquid chromatography tandem mass spectrometry (LC-MS/MS) was used to identify protein substrates and their sites of phosphorylation.

**Figure 2.**
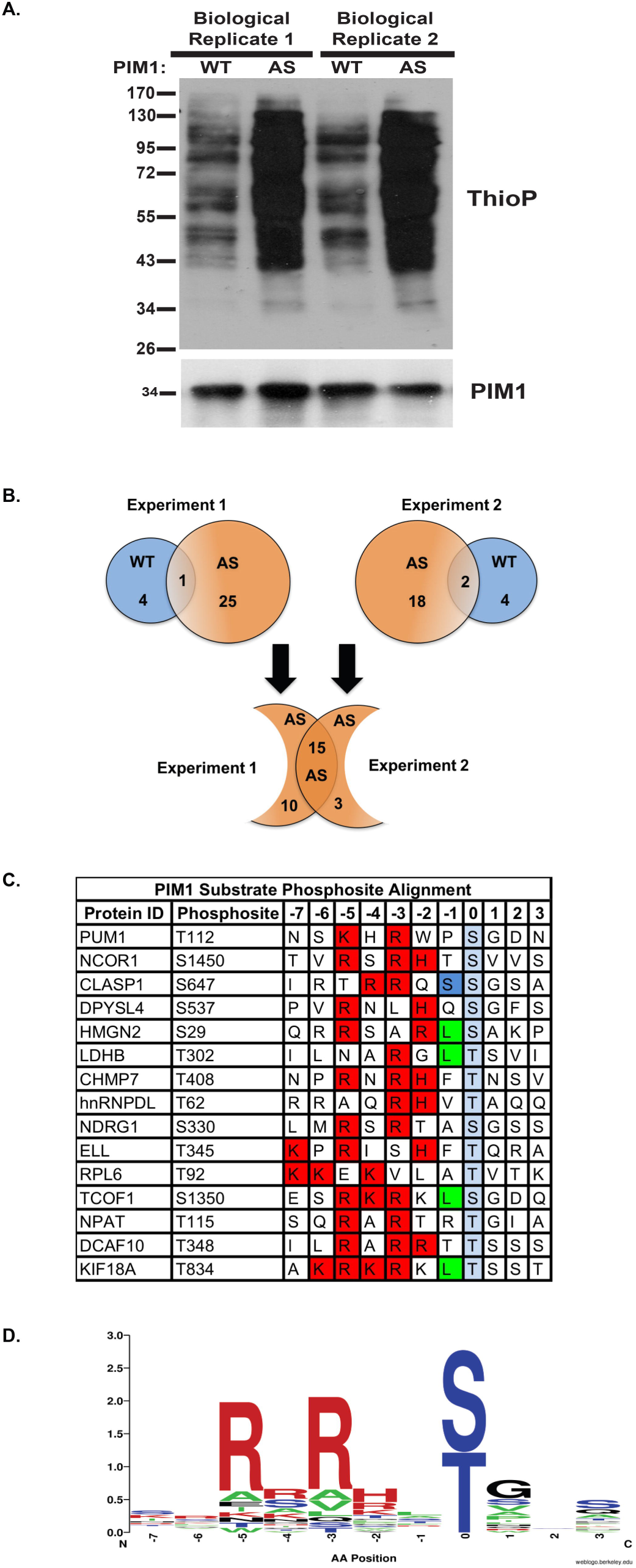
Identification of High-Confidence AS-PIM1 Substrates in Prostate Cancer Cells. A) Western blot of the thiophosphate ester moiety (ThioP) from biological replicates of LNCaP-WT PIM1 and LNCaP-AS PIM1 cell lysates, 0.1% used for the phosphopeptide enrichment. B) Proteins identified from lysates used in Figure 2A from two independent experiments. Blue and orange circles represent proteins identified in WT PIM1 and AS PIM1 expressing LNCaP cells, respectively. C) PIM1 substrate phosphosite alignment. Highlighted residues represent optimal, amino acids based on PIM1 consensus sequence. Red, basic residues. Green, hydrophobic residues. Blue, neutral polar residues. D) Consensus logo generated using substrates in C.

To identify and prioritize high-confidence PIM1 substrates and phosphorylation sites, we used a stringent set of criteria to analyze the mass spectrometry data. Phosphopeptides found in the AS PIM1 datasets, but not in the WT PIM1 control datasets were selected. From a total of 25 protein substrates identified in two biological replicates of AS PIM1 samples (Suppl. Table 2), 15 previously unidentified PIM1 substrates were represented in both replicates (Fig. 2B-C). A PIM1 consensus motif logo was generated from the substrates found in this study (Fig. 2D). This motif conformed to the literature consensus PIM1 phosphorylation motif, R/K-X-R/K-H-X-S/T, where X is a small, neutral amino acid; and an acidic or basic residue (lysine or arginine) at the −3 and −5 positions from the phosphorylation site^30^. Therefore, these proteins likely represent high-confidence PIM1 substrates (Fig. 2C, Suppl. Table 3). We first focused on candidates which strongly adhered to the PIM1 consensus motif, had high peptide counts, and had been previously linked to human prostate cancer. While no previously identified substrates such as AR or c-Myc were found in our screen, we believe our screen identifies the most robust substrates of PIM1 in LNCaP cells. However, we not think that the screen is exhaustive. For example, we previously showed that AR serine 213 is a target of PIM1. We were able to confirm this in LNCaP cells by immunoprecipitating AR from LNCaP expressing AS-PIM1 and show that it can be thiophosphorylated (Fig 1D). Given this result and that the identified substrates conform to the PIM1 consensus phosphorylation motif, we think that the paucity of known PIM1 substrates is not an issue of altered substrate specificity of the AS-PIM1, but rather low abundance of the previously recognized PIM1 substrates, low levels of phosphorylation, or an inability to capture the peptides on MS^21,22^. The combination of our directed thiophosphorylation of AR from LNCaP cells expressing AS-PIM1, coupled with the phosphorylation motif analysis indicating extensive alignment to the PIM1 phosphorylation consensus sequence, gives us high confidence that the substrates we identified represent *bone fide* PIM1 substrates, even if the screen is not saturating.

Several of the PIM1 substrates we identified are involved in the regulation of RNA and DNA metabolism. For example, PUM1, a PUF family RNA-binding protein, regulates translation of sequence-specific genes by binding their 3’ UTRs^31,32^. Substrates NCOR1 and RNA polymerase II elongation factor (ELL) have been implicated in RNA polymerase II pausing and activation, respectively^33,34^. Additional substrates identified in the screen, including CHMP7, TCOF1, NPAT, CLASP1, and KIF18A, have strong associations with mitosis, spindle formation, and microtubule depolymerization^35–40^. These connections to mitosis are intriguing, since PIM1 is known to interact with nuclear mitotic apparatus protein (NuMa) and to promote cell cycle progression during mitosis^41^. Lastly, NDRG1, an established metastasis suppressor involved in cell motility, and has been shown to interact with the WNT receptor LRP6 to block WNT signaling^42^. Taken together, we have identified PIM1 substrates which are involved in regulating cell cycle, nucleic acid metabolism, cell signaling, and cell migration.

To validate the substrates identified in our screen, we tested endogenous thiophosphorylation of PUM1, NDRG1, CHMP7, and KIF18A in LNCaP-AS PIM1 versus LNCaP-WT PIM1 cells. All four proteins were thiophosphorylated in the presence of PhET-ATPγS by AS PIM1 to a greater extent than by WT PIM1 (Fig. 3A-D). To confirm the sites of phosphorylation, we generated serine/threonine to alanine mutants of each substrate and co-expressed them with AS PIM1 in 293T cells. We performed the thiophosphorylation assay, including alkylation with *p*NBM, immunoprecipitated the substrate of interest, and performed Western blot using a thiophosphate ester specific antibody. We observed reduced thiophosphorylation in the phospho-mutant substrates (Fig. 3E-H), indicating that these are the predominant sites of PIM1-mediated phosphorylation for each substrate. This data provides strong evidence that the proteins identified in our screen are bona fide PIM1 substrates in prostate cancer cells.

**Figure 3.**
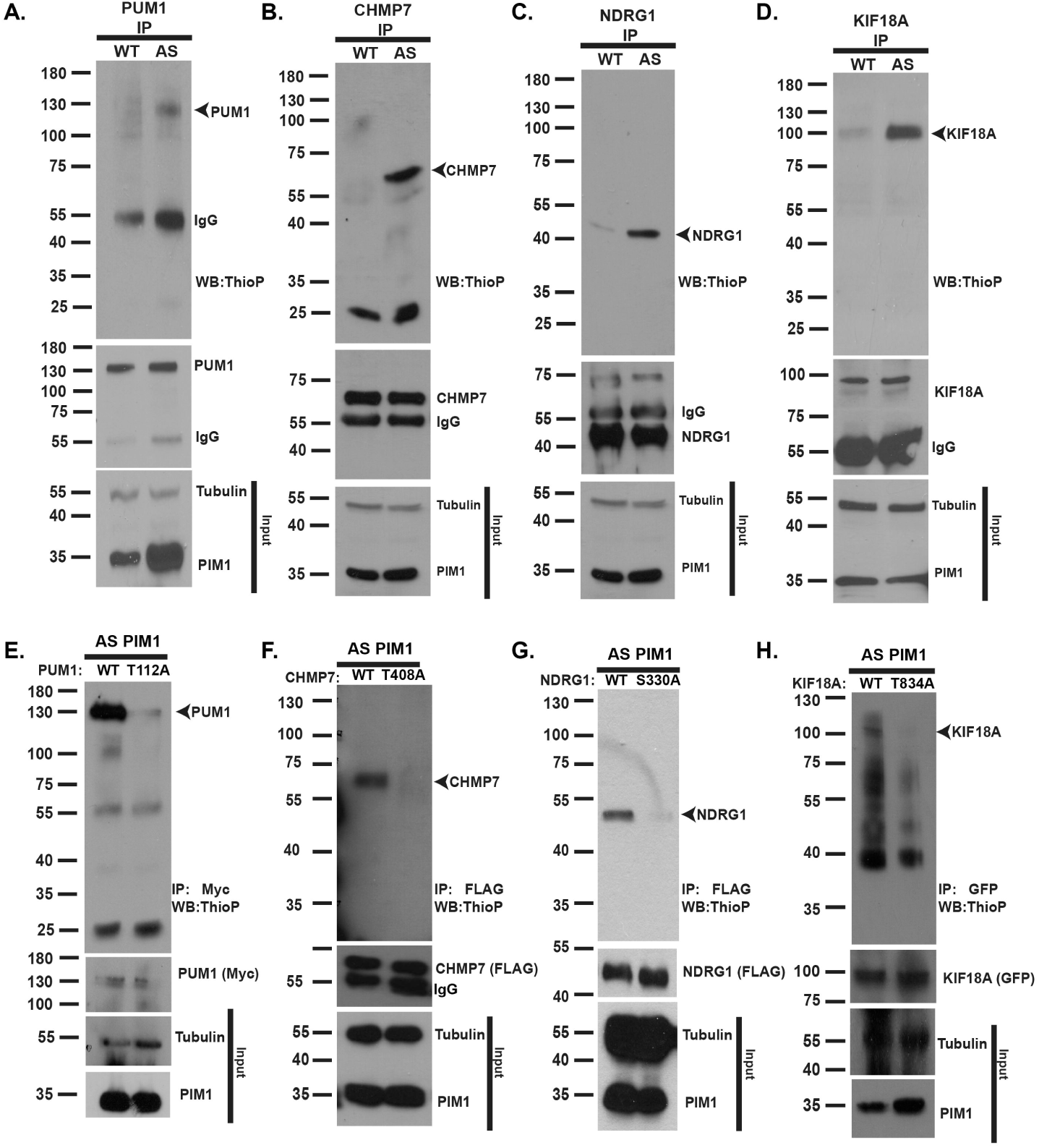
AS PIM1 Directly Phosphorylates Endogenous High-Confidence Substrates in Prostate Cancer Cells. A–D) Validation of endogenous PIM1 substrates. LNCaP-AS-PIM1 thiophosphorylates PUM1 (A), CHMP7 (B), NDRG1 (C), and KIF18A (D). Endogenous proteins were immunoprecipitated from LNCaP-WT-PIM1 and LNCaP-AS-PIM1 and analyzed by Western blot for the presence of thiophosphorylation (ThioP), or immunoprecipitated PUM1, CHMP7, NDRG1, and KIF18A. E-H) Phosphorylation site validation using WT and phosphorylation site mutant substrates. Substrates (Myc-PUM1, FLAG-CHMP7, FLAG-NDRG1, and GFP-6HIS-KIF18A; either wild type (WT) or the indicated PIM1 phosphorylation site mutation) were expressed in 293T cells with AS-PIM1 and thiophosphorylation labeling completed. Substrates were immunoprecipitated using Myc, FLAG, or GFP-magnetic beads, and Western blot performed to detect thiophosphorylated, or the immunoprecipitated substrates. Western blots are representative of two independent experiments.

### NDRG1 phosphorylation varies in prostate cancer cells and tissues

We focused on characterizing the effect of PIM1-dependent serine 330 phosphorylation (pS330) of NDRG1, given that a phosphoproteomic analyses of prostate cancer tissues by Drake *et al*. demonstrated that levels of NDRG1 pS330 were found to be 7.7 times higher in metastatic lesions than in localized prostate cancer^43^ (Sup Table 5). Although several studies have implicated NDRG1 in prostate cancer^44,45^, a link between PIM1 and NDRG1 has not been established. NDRG1 is a member of the alpha/beta hydrolase superfamily, but does not possess catalytic activity^46^. It has been well documented as a potent inhibitor of tumor growth and cancer cell proliferation and an inhibitor of cell migration and invasion^47^. Its expression alters the levels and membrane localization of beta-catenin and the membrane glycoprotein KAI1 (CD82), leading to increased cell-cell adhesion^48^. Additionally, overexpression of NDRG1 reduces metastasis in a rodent model of prostate cancer^49^. In contrast, prostate cancer cells with low NDRG1 expression display increased motility and invasiveness^45^, and reduced expression of NDRG1 was associated with poor overall survival in a variety of cancers, including prostate^50^. NDRG1 is transcriptionally regulated by the androgen receptor^51^. Considering this information, we examined the protein levels of NDRG1 pS330, total NDRG1, and PIM1 by immunohistochemistry using tissue microarrays containing primary prostate tumors and adjacent non-cancer tissue (Suppl. Table 4A-C). Expression of NDRG1 pS330 and AR pS213, another PIM1 substrate, were much more prevalent in specimens with abundant PIM1 (Fig. 4A). Spearman correlation analysis revealed that NDRG1 pS330 levels in prostate cancer are associated with AR pS213 (*r*= 0.3664, *p*=0.01) and PIM1 protein levels (*r*= 0.5038, *p*=0.00003) (Suppl. Table 4C). NDRG1 pS330 was increased in cases with high Gleason grade (Gleason 4/5) compared to tumors with low Gleason grade (Gleason 3) or non-cancer tissue (Fig. 4B, *p*=0.044). This was observed without a significant change in total NDRG1 protein abundance (*p*=0.583), suggesting that the change in NDRG1 pS330 was a result of increased phosphorylation by PIM1 (Fig. 4C).

**Figure 4.**
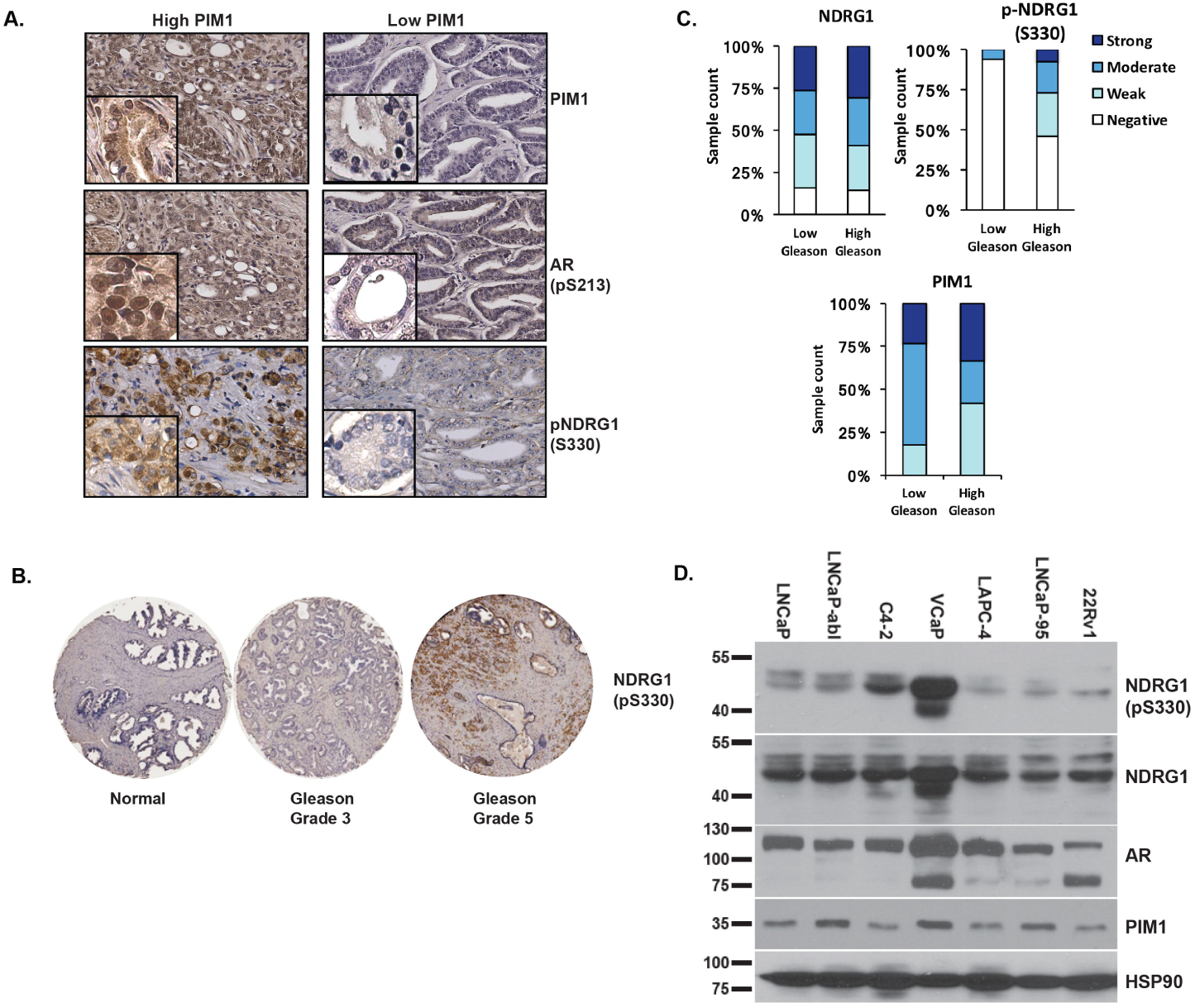
NDRG1 Phosphorylation is Positively Associated with PIM1 Expression and High Gleason Grade in Primary Prostate Cancer. A) Immunohistochemistry (IHC) of PIM1, AR pS213, and NDRG1 pS330 in primary prostate cancer samples (n=67) with low PIM1 and high PIM1 levels. B) IHC of NDRG1 pS330 in normal prostate, low Gleason-grade, and high Gleason cases. C) IHC analysis comparing NDRG1 pS330 and total NDRG1 in low Gleason tumors versus high Gleason tumors (n=76). D) Western blot analysis of NDRG1 pS330, total NDRG1, AR, PIM1 in different prostate cancer cell lines. HSP90 was used as loading control.

We next probed a panel of prostate cancer cell lines for NDRG1 pS330, total NDRG1, and PIM1 by Western blot. AR expressing prostate cancer lines displayed varying levels of NDRG1 pS330. VCaP cells, a prostate cancer cell line with AR amplification, displayed the highest level of total NDRG1 and NDRG1 pS330, consistent with our findings that NDRG1 expression and phosphorylation are androgen regulated (Fig. 4D; Suppl. Fig. 1). Together, these results demonstrate that NDRG1 pS330 is correlated with high grade prostate cancers (high Gleason score), and that NDRG1 pS330 is present in multiple prostate cancer cell lines.

### NDRG1 protein is destabilized by PIM1-mediated phosphorylation

We next investigated the effect of androgens on NDRG1 pS330. As expected, in the absence of androgens, LNCaP cells overexpressing WT PIM^14^ have increased levels of NDRG1 pS330 (Fig. 5A). Treatment with the androgen dihydrotestosterone (DHT) increased NDRG1 pS330 compared to untreated cells. LNCaP cells treated with DHT and with SGI-1776, a potent and specific PIM1 inhibitor^52^, resulted in decreased NDRG1 pS330 (Fig. 5B). PIM1 knockdown by siRNA treatment also reduced NDRG1 pS330 (Fig. 5C). This indicates that NDRG1 phosphorylation at S330 is mediated by PIM1, and inhibition or knockdown of PIM kinase activity decreases phosphorylation.

**Figure 5.**
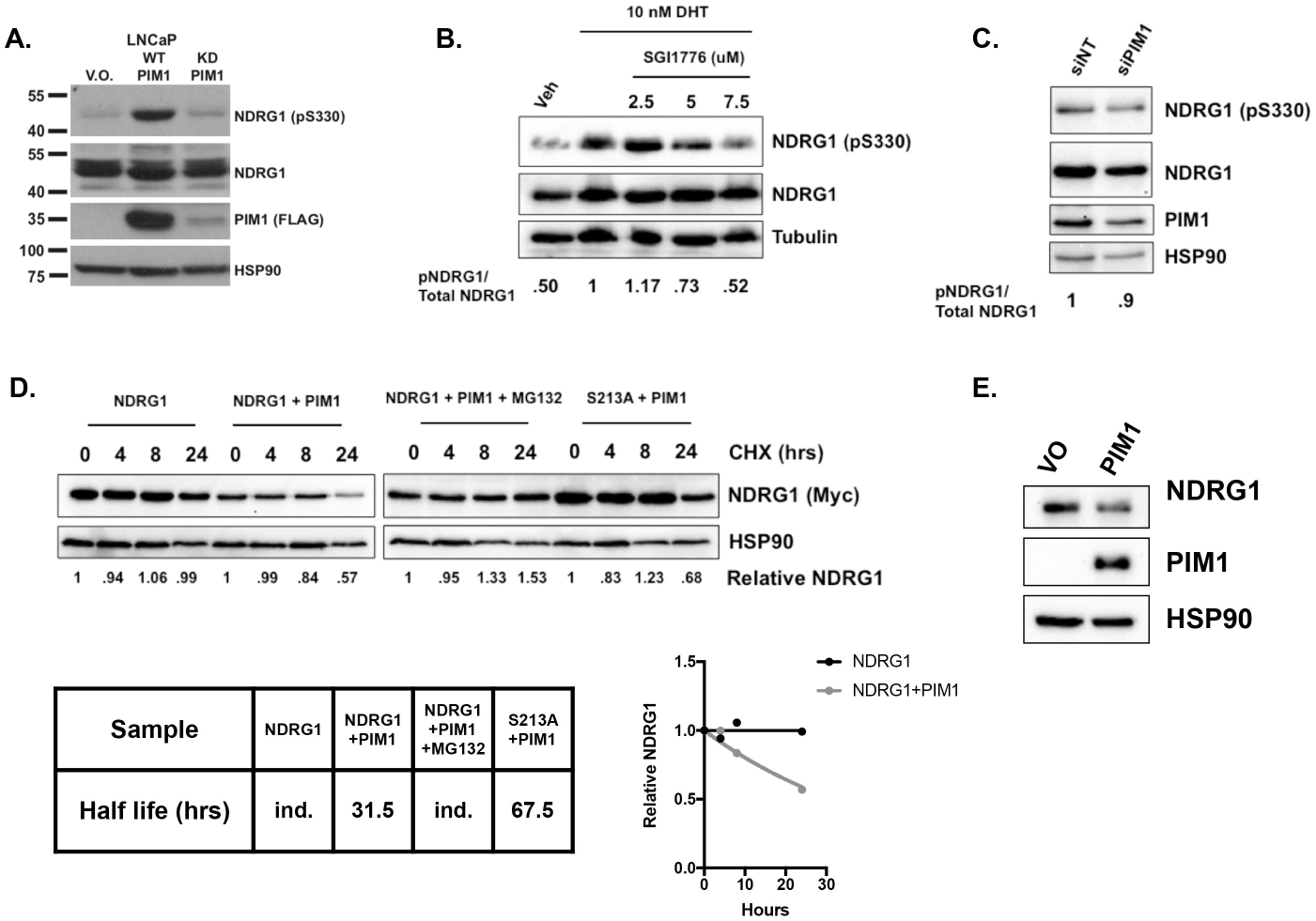
PIM1-mediated NDRG1 pS330 is Androgen-dependent, and Affects Its Stability. A) LNCaP cells stably expressing vector only (LNCaP V.O.), WT PIM1 (LNCaP WT PIM1), or kinase-dead PIM1 (LNCaP KD PIM1) were steroid-starved for 48 h, and Western blot was performed for NDRG1 pS330, total NDRG1, PIM1 (FLAG), and HSP90 (loading control). B) LNCaP and VCaP cells were steroid-starved for 48 h, treated with 10 nM DHT, along with indicated amounts of PIM1 kinase inhibitor SGI-1776 for 24 hours, and Western blot was performed for NDRG1 pS330, total NDRG1, and HSP90. The ratio of NDRG1 pS330/total NDRG1 was quantified using ImageJ software. C) Under the same conditions as Fig. 2B, PIM1 was knocked down using siRNA and Western blot was performed for NDRG1 pS330, total NDRG1, PIM1, and HSP90. Ratios were quantified using ImageJ software. D) Effect of PIM1-dependent phosphorylation on NDRG1 stability. NDRG1 WT and S330A were expressed in 293 cells, with and without PIM1. Cells were treated with cycloheximide for the indicated times, lysates prepared and blotted for NDRG1 and HSP90. Additionally, WT NDRG1+PIM1 was treated with MG-132. Extrapolated half-lives of NDRG1 under the various conditions are shown. E) PIM1 overexpression in LNCaP cells results in loss of endogenous NDRG1.

To determine whether phosphorylation affects the stability of NDRG1, we examined NDRG1 stability by blocking protein synthesis with cycloheximide and tracking NDRG1 abundance over time. We co-expressed WT or phospho-mutant (S330A) NDRG1 with either empty vector or WT PIM1 in 293 cells. Compared with empty vector alone, co-expression of WT NDRG1 with WT PIM1 led to a significant reduction in NDRG1 protein stability (Fig. 5D). NDRG1 is exceedingly stable in the absence of PIM1, with its abundance unchanged over 24 hours in the presence of cycloheximide. However, the addition of PIM1 resulted in a decrease in NDRG1 abundance by 50% at 24 hours (Fig 5D). Treatment with the proteasome inhibitor MG-132 in the presence of PIM1 restored NDRG1 stability, indicating that degradation is occurring via the proteasome. By contrast, NDRG1 S330A mutant was resistant to degradation in the presence of PIM1 (Fig 5D). Finally, we examined the endogenous levels of NDRG1 in LNCaP cells in the presence of PIM1 and found that PIM1 overexpression results in a reduction of NDRG1 protein (Fig 5E). Collectively, we find that PIM1 expression leads to increased levels of NDRG1 pS330, and PIM1-dependent NDRG1 phosphorylation decreases NDRG1 protein stability.

### Non-phosphorylated NDRG1 localizes to the nucleus and interacts with the androgen receptor

We next examined the subcellular distribution of NDRG1 pS330 and total NDRG1 in prostate cancer cells. PIM1 has been reported to be mainly localized to the cytoplasm^53^. LNCaP and VCaP cells were separated into cytoplasmic, membrane, soluble nuclear, and chromatin bound fractions, and probed for NDRG1 pS330 and total NDRG1 by Western blot. In both cell lines, NDRG1 pS330 was largely detected in the cytoplasmic fraction, with a small amount of NDRG1 pS330 in the membrane fraction of VCaP cells. The faster migrating NDRG1 species in the soluble nuclear fraction from LNCaP cells may represent a previously described C-terminally truncated variant of NDRG1^54^. By contrast, total NDRG1 was detected across all fractions, including the nuclear fractions (Fig. 6A). We hypothesized that phosphorylation may prevent NDRG1 from accumulating in the nucleus.

**Figure 6.**
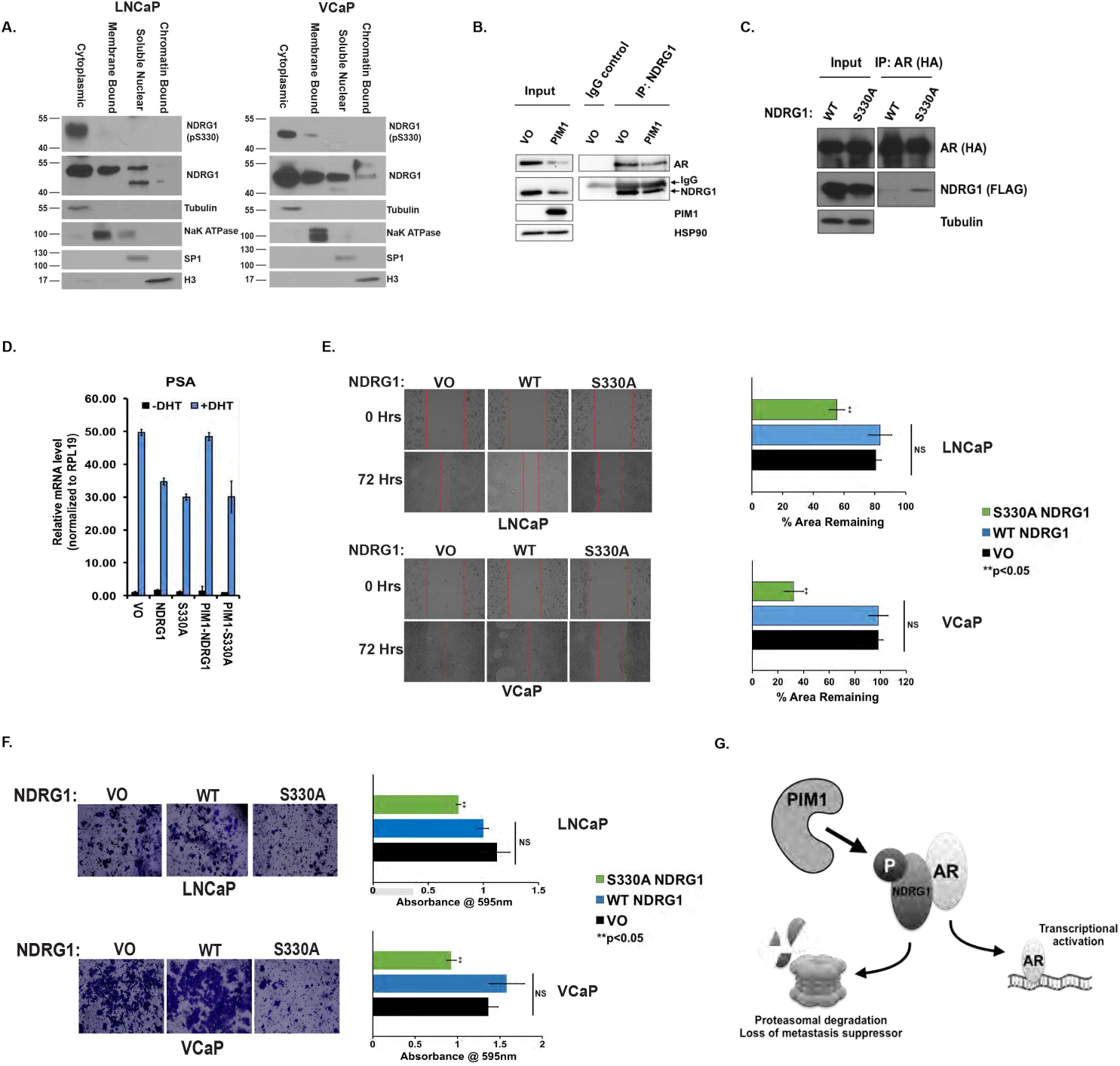
NDRG1-AR interaction is modulated by NDRG1 phosphorylation. A) LNCaP and VCaP cells were fractionated into membrane, cytoplasmic, soluble nuclear and chromatin bound fractions and blotted for NDRG1 pS330 and NDRG1. Markers for each compartment were also blotted for to ensure the fidelity of the fractionation. B) Endogenous NDRG1 co-immunoprecipitates with AR in LNCaP cells. C) AR interacts more robustly with non-phosphorylated NDRG1. 293T cells were transfected with either NDRG1 WT or NDRG1 S330A mutant (FLAG-tagged) and WT-AR (HA-tagged). AR was immunoprecipitated using HA beads, and Western blot performed for AR (HA) and NDRG1 (FLAG). D) LNCaP transfected with indicated genes were steroid-starved for 48 h, and subsequently treated with 10 nM DHT. mRNA levels for PSA were measured relative to RPL19 by qRT-PCR. The experiment performed in triplicate and error bars represent the standard deviation. E-F) Migration and matrigel invasion assays. LNCaP-PIM1 and VCaP-PIM1 expressing cells were transfected with vector only (VO), NDRG1 WT, or NDRG1S330A. Cell migration was determined using a scratch assay. Micrographs of the cells at 0 and 72 h are shown, and percent area remaining after 72 h was quantitated. Invasion assay through matrigel was performed in LNCaP-PIM1 and VCaP-PIM1 cells expressing the indicated NDRG1 constructs, and cells invading through matrigel were stained and shown as micrographs. The stain was extracted from the cells that migrated through the matrigel and quantitated spectrophotometrically at 595 nm wavelength. G) Model illustrating the effect of PIM1 phosphorylation on NDRG1 and its co-repression of AR.

Prostate cancer is highly dependent on AR transcriptional activity^55^. For this reason, we examined whether NDRG1 interacts with AR and affect its transcriptional activity. We immunoprecipitated endogenous NDRG1 from LNCaP cells, with or without PIM1 overexpression (Fig. 6B). We were able to observe co-immunoprecipitation of AR with NDRG1. Additionally, consistent with our previous results, the overexpression of PIM1 reduced the abundance of NDRG1 protein, resulting in a reduction in immunoprecipitated NDRG1 and AR. Next, to determine whether S330 phosphorylation affects the interaction between NDRG1 and AR, we over-expressed WT or S330A NDRG1 in 293T cells (Fig. 6C). We immunoprecipitated AR and observed that it interacts more robustly with NDRG1 S330A than WT NDRG1. This suggests that non-phosphorylated form of NDRG1 is more capable of interacting with AR potentially through alterations in its abundance or localization.

Because we observed an interaction between AR and NDRG1, we tested whether NDRG1 affected AR transcriptional activity. We transfected LNCaP cells with NDRG1 or the non-phosphorylatable NDRG1 S330A, in the absence and presence of PIM1, and assayed the expression of AR target genes PSA and NKX3.1. Expression of WT NDRG1, and to a greater extent S330A, decreased androgen-dependent expression of PSA and NKX3.1 (Fig. 6D, Suppl. Fig. 2A). Co-expression of PIM1 with NDRG1 rescued the PSA and NKX3.1 expression in the presence of WT NDRG1, but not NDRG1 S330A. This suggests that NDRG1 negatively regulates AR activity, and that PIM1-dependent NDRG1 pS330 reverses this effect. Collectively, this data suggests that non-phosphorylated NDRG1 interacts with and represses AR transcriptional activity, and upon phosphorylation at S330 by PIM1, this interaction and co-repression is reduced.

### NDRG1 pS330 is associated with inhibition of prostate cancer cell migration and invasion

Multiple reports have illustrated the role of NDRG1 as a metastasis suppressor^42,45,56^. For example, overexpression of NDRG1 in a mouse xenograft model reduced lung and liver metastasis^57^. NDRG1 regulates the expression of the membrane protein KAI1 (CD82), which interacts with adjacent cells via the membrane protein gp-Fγ to inhibit metastasis^57^. Moreover, NDRG1 expression in HepG2 cells leads to increased membrane-localized beta-catenin, which promotes a less migratory phenotype and greater cell-cell contact^45^. In light of these findings, and given the previously demonstrated ability of PIM1 to promote cell migration and invasion^58,59^, we investigated the effect of NDRG1 phosphorylation on the migration and invasion of prostate cancer cells. To examine migration, we exogenously expressed WT NDRG1 or NDRG1 S330A in PIM1 overexpressing LNCaP and VCaP cells (Suppl. Fig. 2C), and used a scratch assay to monitor migration over 72 hours. In LNCaP-WT PIM1 expressing cells, there was an 80% scratch area recovery after 72 hours. WT NDRG1 overexpression was unable to suppress this cell migration, but in cells expressing NDRG1 S330A, there was only a 50% recovery (Fig. 6E). Likewise, in VCaP cells expressing WT PIM1 alone or with WT NDRG1, 100% scratch closure was observed after 72 hours, and the area recovery was only 33% in cells expressing NDRG1 S330A (Fig. 6E). Knockdown of NDRG1 by siRNA in LNCaP cells also increased cell migration (Suppl. Fig. 2B). Similar results were observed in a matrigel invasion assay. In LNCaP and VCaP cells co-expressing WT PIM1 and WT NDRG1, there was no difference in invasion. However, in cells expressing NDRG1 S330A, there was a significant reduction in cell invasion (Fig. 6F). KAI1 expression is also increased in with overexpression of NDRG1 S330A in LNCaP cells (Suppl. Fig. 2D). These results suggest that phosphorylation of NDRG1 reduces its function as a suppressor of migration and invasion, and that blocking PIM1-dependent NDRG1 phosphorylation could restore its metastasis suppressor function

## DISCUSSION

Our screen is the first to use an unbiased, direct method to identify PIM1 substrates in prostate cancer cells. We identified 25 previously unknown PIM1 substrates and their phosphorylation sites. We demonstrated that the specificity of the AS PIM1 was retained, as AS PIM1 is still able to thiophosphorylate the known PIM1 substrate, AR^14^. Of the 25 substrates that were present in AS PIM1 and not WT PIM1, 15 were identified in two biological replicates, and the phosphorylation sites conform to the PIM1 consensus sequence. We validated four of these targets using alkylation and detection by western blot. Our screen, however, was not comprehensive, and did not identify other previously identified PIM1 substrates, such as c-MYC. This may reflect low abundance of the phosphorylated substrates in LNCaP cells, or may be due to the fact that not all tryptic peptides are equally eluted from iodoacetyl beads or equally detectable by mass spectrometry. Additionally, our approach represents a direct, unbiased approach to identifying PIM1 substrates in prostate cancer cells. Therefore, these substrates may be the most representative of PIM1 activity in prostate cancer. In fact, PIM1 has overlapping substrate specificity with AKT protein kinases: for example, both have been shown to phosphorylate the pro-apoptotic protein BAD^60,61^. However, data from *in vivo* studies using knock-out mice indicate that AKT and PIM kinases are mechanistically distinct in the control of hematopoietic cell proliferation^62^. This reinforces the notion that identification of direct PIM1 targets is key to elucidating its function in a given cell type.

We have found that PIM1 substrates are involved in multiple physiological processes. Network enrichment analysis of all 25 substrates using Metascape^63^ [http://metascape.org] revealed functional classes including chromosome segregation, pre-RNA complex, nucleotide catabolic processes, transcriptional regulation, and infectious disease (Suppl. Fig. 3A). This is consistent with studies of PIM1’s effect on cell cycle, transcription, and cell motility. In addition, a majority of the substrates fall into a protein interaction network that suggests PIM1 phosphorylation may coordinate protein complexes to impact cellular function (Suppl. Fig. 3B). Further studies will be required to validate the interaction network and determine whether these associations are phosphorylation dependent.

Among the newly identified PIM1 substrates, we focused on NDRG1. NDRG1 was shown to be as an inhibitor of metastasis. Its expression alters the expression and membrane localization of beta-catenin and the membrane glycoprotein KAI1 (CD82), leading to increased cell-cell adhesion^57^. NDRG1 is transcriptionally regulated by AR, which is also phosphorylated by PIM1. In addition, Drake *et al*. reported that the level of NDRG1 pS330 was higher in metastatic castration resistant prostate cancer (CRPC) lesions than in treatment naïve, localized prostate cancer^43^. This was the only phosphorylation site identified in our screen that was increased in metastatic compared to localized prostate cancer (Suppl. Table 5). Immunohistochemical analysis of prostate cancer tissue microarrays showed a significant increase in NDRG1 pS330 in malignant compared to benign tissues. This correlates with the low expression of PIM1 in benign tissues and moderate to high PIM1 expression in prostate cancer^12^. PIM1-dependent phosphorylation of NDRG1 appears to also reduce its stability, with the NDRG1 S330A mutant showing an increased half-life in the presence of PIM1. Previous studies have suggested that NDRG1 protein stability is reduced by the fusion of SUMO2 to NDRG1^64^. Whether phosphorylation of NDRG1 results in NDRG1 sumolyation to control protein stability remains to be addressed.

The migratory and invasive potential of cancer cells are key mediators of metastasis^18^. PIM1 expression has been correlated with lymph node metastasis and poor prognosis in patients with lung adenocarcinoma and squamous cell carcinoma^65^. We were interested in how PIM1 kinase activity could promote metastasis via one of the substrates we identified. Indeed, the non-phosphorylatable NDRG1 S330A suppressed migration and invasion of LNCaP and VCaP cells. We have shown that PIM1 phosphorylation of NDRG1 reduces its stability, reduces its nuclear localization, reduces its interaction with AR, and therefore relieves AR co-repression. This suggests that PIM1-mediated phosphorylation of NDRG1 inhibits its ability to exert metastasis-suppressive functions in prostate cancer. Our results establish an understanding of how phosphorylation inhibits the function of NDRG1 in suppressing migration and invasion. Based on these findings, we propose a model whereby PIM1 is, in part, imposing its oncogenic function by phosphorylating NDRG1 to reduce its metastasis suppressor activity (Fig. 6G). Thus, identifying direct substrates has allowed for the characterization of the mechanism by which PIM1 mediates an oncogenic phenotype – namely cell migration and invasion - in the context of prostate cancer. We anticipate that characterization of additional PIM1 substrates will lend further mechanistic insights into the oncogenic properties imparted by PIM1 in prostate cancer.

## Methods

### Cell Culture and Reagents

Cell lines used in this study were purchased from ATCC, with the exception of LNCaP-95^66^, LNCaP-abl^67^, and LAPC4 cells, which were gifts from Dr. J. Luo (Johns Hopkins University), Dr. Z. Culig (Innsbruck Medical University, Austria), and Dr. R. Reiter (University of California, Los Angeles) respectively. C4-2 cells were purchased from Characterized Cell Line Core Facility at MD Anderson Cancer Center (Houston, TX). Both LNCaP-95 and LNCaP-abl cells have been authenticated by short tandem repeat profiling. 293T and VCaP cells were cultured in DMEM supplemented with 10% fetal bovine serum, 1% penicillin/ streptomycin. LNCaP, 22Rv1, and C4-2 cells were cultured in RPMI (containing phenol red and L-glutamine) supplemented with 10% fetal bovine serum, 1% penicillin/ streptomycin. LNCaP-abl and LNCaP-95 cells were cultured in RPMI (without phenol red) supplemented with 10% charcoal-stripped fetal bovine serum, 1% penicillin/ streptomycin, 1% L-glutamine. LAPC-4 cells were cultured in IMDM supplemented with 10% fetal bovine serum, 1% penicillin/ streptomycin. All cell lines were regularly assessed for mycoplasma contamination.

### Plasmids, Primers, and Antibodies

Constructs utilized are as follows: pCDNA 3.1 (+)-AR-FLAG^14^, NDRG1 (HG14119-CF, Sino Biologicals), CHMP7 (HG14273-CF, Sino Biologicals), PUM1 (RC201219, Origene), KIF18A (Addgene plasmid # 23002), pCDNA 3.1 (+)-WT PIM1^14^, pCDNA 3.1 (+)-KD (K67M)^14^, pLB(N)CX-WT PIM1^14^, pLB(N)CX-KD PIM1^14^. pCDNA 3.1 (+)- AS (L120G) PIM1 and pLB(N)CX-AS PIM1 (L120G) was generated using site-directed mutagenesis. WT, KD, and AS PIM1 plasmids contain the 33kDa PIM1-S isoform. Primers are as follows for mutagenesis: PIM1 L120G (5’-gctcgggcctctccccgatcaggacgaaac-3’), NDRG1 S330A (5’-cgctggaaccagcggctgtgcggga-3’), PUM1 T112A (5’-gcaaacatcgatggcctgctggggataacattcat-3’), KIF18A T834A (5’-actgtttgatgtagaacttgctaatttccgtttccttttggcag-3’), CHMP7 T408A (5’-cacgctgttggcaaaatgcctattgcgggggt-3’). Resulting products was transformed in competent DH5α cells (Thermofisher). Site-directed mutagenesis was performed using the QuikChange Lightning Multi Site-Directed Mutagenesis kit (210515, Agilent) according to manufacturer’s instructions. All plasmids were sequenced to confirm the mutations and to ensure no additional changes in the sequence were introduced. The antibodies used were as follows: PIM1 (sc-13513; Santa Cruz Biotechnology), NDRG1 (HPA006881, Sigma), NDRG1 pS330 (ab124713, Abcam), AR (sc-7305, Santa Cruz Biotechnology), HSP90 (610418, BD Biosciences), FLAG (F3165, Sigma), α-Tubulin (ab7291, Abcam), CHMP7 (HPA036119, Sigma), Thiophosphate ester antibody [51-8] (ab92570, Abcam), KIF18A (C-19, Santa Cruz Biotechnology), PUM1 (ab92545, Abcam), NaK^+^ ATPase (EP1845Y, Abcam), SP1 (PA5-29165, Thermofisher), Histone H3 (ab1791, Abcam).

### Quantitative RT-PCR

Total RNA was extracted and quantitative RT-PCR was performed as previously described ^14^. Primers can be found in Supplemental Table 6.

### Transfection and Transduction

Transient transfections were carried out utilizing Lipofectamine 2000 (Invitrogen) according to the manufacturer’s protocol. Viral transductions were carried out in 293T cells (4.5 x 10^6^ per 10cm plate) utilizing 5 μg target retroviral construct, pLB(N)CX-ORF [where ORF represents the gene of interest], 2μg VSV-G (Addgene plasmid # 8454), and 3μg pGag-POL (Addgene plasmid # 35614) utilizing Lipofectamine (Invitrogen). Target cells, plated at 2 x 10^6^ per 6 cm plate, were treated with viral supernatant, supplemented with polybrene (8 μg/mL, H9268, Sigma dissolved in dH_2_O) for 4 hours, and media replaced. Viral supernatant treatment completed twice over a 48 h period, and pools selected using Blasticidin S (A1113903, Thermofisher) (8μg/mL). Selected pools were maintained in Blasticidin S (1μg/mL).

### Thiophosphorylation of PIM1 Substrates

Thiophosphorylation experiments were carried out as described with minor modifications^27^. Briefly, 2.0 x 10^5^ HEK293T cells were transiently transfected with 1μg of WT or AS (L120G) PIM1 and 1μg of candidate substrate in poly-d-lysine coated 6-well plates using Lipofectamine 2000 according to manufacturer’s protocol. After 48 h cells are washed 2X PBS, and treated with 150 μL thiophosphorylation buffer (20 mM HEPES pH 7.3, 100 mM KOAc, 5 mM NaOAc, 2 mM MgOAc_2_, 1 mM EGTA, 10 mM MgCl2, 0.5 mM DTT, 5 mM creatine phosphate (#2380; Calbiochem), 57 mg/ml creatine kinase (#2384; Calbiochem), 30 mg/ml digitonin (D141, Sigma), 5 mM GTP (G8877, Sigma), 0.1 mM ATP (A1852, Sigma), 0.1 mM Adenosine-5’-O-(3-thiotriphosphate) (ATPγS), or 0.1 mM N^6^-Phenyladenosine-5’-O-(3-thiotriphosphate) (Phe-ATPγS), or N^6^-(2-Phenylethyl)adenosine-5’-O-(3-thiotriphosphate) (PheET-ATPγS) (# A060, P044, P026, respectively, Biolog, Germany), 0.45 mM AMP (A1752, Sigma), 10 mM calyculin A (#990; Cell Signaling), and 1X protease inhibitor cocktail (#5871, Cell Signaling) for 20 min with gentle rotation at room temperature. 150μL of a modified 2X RIPA buffer (100 mM Tris [pH 8.0, 300 mM NaCl, 2% NP-40, 0.2% SDS, and 20 mM EDTA) containing 5 mM *p*NBM (alkylating agent) was added and incubated at room temperature for 60 min. Following alkylation, lysates were collected and cell debris removed by centrifugation at 14,000 RPM for 10 min at 4°C. Protein lysates were suspended in 6X SDS loading buffer and boiled for 5 min. Proteins were resolved by SDS-PAGE and analyzed with the designated antibodies.

For mass spectrometric analysis, 4.0 x 10^6^ LNCAP cells stably expressing either WT-PIM1 or AS (L120G) PIM1 were plated in poly-d-lysine coated 100 mm plates and allowed to attach overnight. A total of 10 plates per replicate were utilized for mass spectrometric analysis. For each plate, 500 μL of thiophosphorylation buffer was used. Protein samples (3 mg) were spiked with 1 μg of thiophosphorylated myelin basic protein (MBP) as enrichment control and the samples were lysed and digested into peptides as described in Hertz et al^27^ with the following modifications. Briefly, 3 mL of cell lysate samples were resuspended in 9 mL of denaturation buffer (8M urea solution in 100mM ammonium bicarbonate, 2 mM EDTA, 1M TCEP). Samples were reduced at 55° C for 1 h. Post reduction samples were cooled to room temperature and diluted using 50mM ammonium bicarbonate so that the final urea concentration was 2M to facilitate enzymatic digestion. To ensure the samples remained in a reduced state, additional TCEP was added to a final concentration of 10mM. Samples were digested overnight at 37° C using sequencing grade modified trypsin. Samples were acidified using trifluoroacetic acid (TFA) to a final concentration of 0.2% to stop digestion. Peptides were loaded onto a C18 Sep-Pak cartridge (Waters) equilibrated with 0.1% TFA and desalted by washing with 9 mL of 0.1% TFA. Peptides were eluted stepwise with 40% followed by 80% acetonitrile in 0.5% acetic acid and the eluatates were combined and concentrated using a SpeedVac. The desalted peptide mixture was subjected to covalent capture of thiophosphorylated peptides as described in^27^.

Using the auto sampler of an Easy nLC1000 (Thermo Scientific), an aliquot of each enriched sample was injected onto a trap column (Acclaim® PepMap 100 pre-column, 75 μm × 2 cm, 3 μm, 100 Å, Thermo Scientific) coupled to an analytical column (EASY-Spray PepMap column, 50 × m × 75 μm ID, 2 μm, 100 Å, Thermo Scientific). Peptides were gradient eluted into a Q Exactive (Thermo Scientific) mass spectrometer using a 60 min gradient from 2% to 31% solvent B (90% acetonitrile, 0.5% acetic acid), followed by 10 minutes from 31% to 40% solvent B and 10 minutes from 40% to 100% solvent B. The Q Exactive mass spectrometer acquired high resolution full MS spectra with a resolution of 70,000 (at *m/z* 400), an AGC target of 1e6 with maximum ion time of 120 ms, and a scan range of 400 to 1500 *m/z*. Following each full MS, 20 data-dependent high resolution HCD MS/MS spectra were acquired using a resolution of 17,500 (at *m/z* 400), AGC target of 5e4 with maximum ion time of 120 ms, one microscan, 2 *m/z*, isolation window, fixed first mass of 150 *m/z*, normalized collision energy (NCE) of 27, and dynamic exclusion of 30 seconds. The MS/MS spectra were searched against the Uniprot human reference database using Byonic ^68^ (Protein Metrics) with the following parameters: oxidation of methionine (M), deamidation of asparagine (N) and glutamine (Q), phosphorylation of serine (S), threonine (T) and tyrosine (Y) were selected as variable modifications. Both precursor and fragment mass tolerances were set to 10 ppm. All identified peptides were filtered using a Byonic score of >200. And a false-discovery rate (FDR) of 0.01 was applied for protein level identification.

### Immunoprecipitation of Candidate Substrates

For immunoprecipitation after thiophosphorylation and alkylation referred to in “Thiophosphorylation of PIM1 Substrates” (Fig. 3), 40 μl out of 500 μl lysate was retained to evaluate expression. The remaining lysate was incubated with 25 μL of FLAG agarose beads (M2, Sigma, F2426) pre-washed 3 X in bead wash buffer (50 mM Tris HCl pH 7.5, 100 mM NaCl, 5 mM EDTA, and 0.4% Triton-X), and incubated for 2 h at 4°C with end-over-end rotation. In order to recover immunoprecipitated protein, beads were washed 5X with bead wash buffer containing protease inhibitors. To elute proteins, beads were incubated with 100 μL TBS containing FLAG peptide for 1 h at 4°C with end-over-end rotation. Beads were pelleted and the 6X SDS loading buffer added to eluted proteins in the supernatant and boiled for 5 min. Proteins were resolved by SDS-PAGE and analyzed by western blotting with the appropriate antibodies.

### NDRG1 Protein Stability Assay

293 cells were transfected with indicated constructs using Lipofectamine 2000 (Invitrogen) according to manufacturer’s protocol. Eighteen hours post-transfection, 50 μg/mL cycloheximide (C7698, Sigma) was added to cells for the indicated times. Cells were lysed in 1X RIPA supplemented with 1X complete protease inhibitors, and 10 μM calyculin A. Protein lysates were resuspended in 6X SDS loading buffer and boiled for 5 min. Proteins resolved by SDS-PAGE and analyzed with NDRG1 and PIM1 antibodies. For LNCaP cells expressing vector only or PIM1^14^, cells were treated with cyclohexmide for the indicated times, lysates prepared and NDRG1 and PIM1 abundance determined. For MG132 experiments, cells were treated with 10 μM MG-132 for the duration of the treatment in the presence of cyclohexmide.

### Migration and Matrigel Invasion Assay

For the migration assay, LNCaP and VCaP cells were plated in 6-well plates, at 500,000 cells per well and transfected with indicated constructs using Lipofectamine 2000 according to manufacturer’s instructions. After 48 h, cells were plated into 2 well silicone insert with a defined cell-free gap, according to manufacturer’s instructions (#81176, Ibidi USA). LNCaP cells were plated at 350,000/mL, and VCaP cells were plated at 850,000/mL. After an overnight incubation, the inserts were removed, and images captured after 72 h. Images were obtained at 5X magnification utilizing an EVOS FLc. The area remaining was measured by ImageJ software in arbitrary units, with error bars representing standard deviation for three independent experiments performed in triplicates (**P value=0.05, Student’s t-test). For the invasion assay, matrigel-coated chambers were rehydrated with serum-free DMEM. 100,000 LNCaP or VCaP cells were added to the upper chamber in 500µL of media. In the lower chamber 500µL of complete media was added to serve as the chemoattractant. After 48 h, cells that did not invade matrigel were cleared away using a cotton swab, and remaining cells were stained with 0.05% crystal violet dye. For quantification, chambers were incubated for 2 mins in 150µL of 10% acetic acid in dH_2_O to solubilize the dye taken up by the cells and quantified spectrophotometrically at 570 nm.

### Human Prostate Cancer Tissue Microarrays, Immunohistochemistry, and Scoring

Human prostate tissue microarrays are from US Biomax, Inc. (Rockville, MD) (cat # PR803a and PR807b). Tissues slides were deparaffinized and rehydrated in xylenes and a series of graded ethanol. Antigen retrieval was performed with 0.01 M citrate buffer at pH 6.0 for 20 minutes at 95°C. Slides were allowed to cool for another 30 minutes, followed by sequential rinsing in phosphate-buffered saline with 0.01% Triton X (PBS-T). Endogenous peroxidase activity was quenched by incubation in PBS-T containing 3% hydrogen peroxide. Each incubation step was carried out at room temperature and was followed by three sequential washes in PBS-T. Sections were incubated in 5% normal goat serum in RT for 1 h before incubation with rabbit polyclonal NDRG1 pS330, total-NDRG1, PIM1, AR pS213 and total AR (441) antibodies, overnight at 4°C. The next day, slides were washed with PBS-T three times and were incubated with biotinylated secondary antibody for 1 h, peroxidase-labeled streptavidin (Vectastain system, Vector Laboratories) for 1 h, and diaminobenzidine substrate for peroxidase-based immunohistochemistry (Cardassian DAB Chromogen, Biocare Medical). Slides were counter-stained with hematoxylin (Vector Laboratories) and dehydrated before mounting. The intensity of IHC staining was scored as negative (0), weak (1), moderate (2), or strong staining (3). Comparison between NDRG1 p330 score and AR pS213 score in high versus low PIM1 expressing cases was analyzed by Student’s t-test. Relationship between NDRG1 p330 in high and low Gleason grade cases was analyzed by Student’s t-test. The association among NDRG1 p330, total-NDRG1, and PIM1 were calculated using Spearman correlation coefficient analysis.

## AUTHOR CONTRIBUTIONS

R.L., S.R., B.U. M.G., and S.L. designed the study. R.L., Y.W., S.R., J.S. and S.N. carried out experiments; R.L., Y.W., B.U., S.R. and S.N. analyzed data and performed statistical analyses. B.U., M.G., and S.L. provided supervisor oversights. R.L., S.R., Y.W., M.G., B.U. and S.L. wrote the manuscript, and all authors approved the manuscript.

## ACKNOWLEDGMENTS

We thank Susan Ha and members of the Logan and Garabedian labs for critically reading the manuscript. This work was supported by grants from the Ford Foundation and the HHMI Gilliam Fellowship (RJL), and NIH R01CA112226 (SKL) and T32CA009161 (SR, JAS). The mass spectrometric experiments where in part supported by NYU School of Medicine and the Laura and Isaac Perlmutter Cancer Center Support grant P30CA016087 from the National Cancer Institute.

---

**Supplemental Figure 1.** NDRG1 expression is stimulated by androgen-treatment in LNCaP cells. LNCaP cells were steroid-starved for 48 h, and treated with 10 nM DHT for 24 h. mRNA levels for PSA, and PIM1 substrates relative to RPL19 were analyzed by qRT-PCR.

**Supplemental Figure 2.** Supplemental Figures associated with Figure 6. A) LNCaP cells were transfected with the empty vector, NDRG1 or NDRG1 S330A, without or with PIM1, steroid-starved for 48 h, and treated with 10 nM DHT for 24 h. mRNA expression of Nkx3.1 was measured by qRT-PCR relative to RPL19. B) LNCaP cells were transfected with either non-targeting siRNA (siControl) or NDRG1 siRNA (siNDRG1). After 48 h, cells were plated for migration assay, and area remaining quantified by ImageJ analysis. Inset illustrates NDRG1 knockdown by siRNA by Western blot. C) Western blot analysis from LNCaP and VCaP cells of exogenously expressing empty vector (-), wild type NDRG1 (WT), or NDRG1 S330A (SA) in the presence of WT PIM1. HSP90 was used as loading control. D) (Pertaining to Figure 6D-E) LNCaP cells were transfected with either NDRG1 WT or NDRG1 S330A and/or WT PIM1 or kinase dead (KD) PIM1, and KAI1 gene expression was measured by qRT-PCR relative to RPL19.

**Supplemental Figure 3.** Metascape analysis of gene ontology and protein-protein interactions of PIM1 substrates. A) Heatmap of enriched terms of the 25 PIM1 substrates from LNCaP cells colored by p-values. B) Protein-protein interaction network of the PIM1 substrates from LNCaP cells.

**Supplemental Table 1.** PIM1 gatekeeper residue alignment with kinases that analog-sensitive mutants have been generated for ^22,28,69–72^.

**Supplemental Table 2.** PIM1 substrates identified from LNCaP cells. UniProt accession ID, phosphorylation site, peptide count from two independent experiments, validation status, and function are listed.

**Supplemental Table 3.** PIM1 substrate phosphorylation site alignment.

**Supplemental Table 4.** Prostate tumor microarray sample data, scoring and statistical analysis.

**Supplemental Table 5.** PIM1 substrate phosphorylation in metastatic versus primary prostate cancer from Drake *el al.^43^*

**Supplemental Table 6.** Primers for qRT-PCR of PIM1 substrates.

