## Supplemental_Figs_Tables for "Identification of PIM1 substrates reveals a role for NDRG1 in prostate cancer cellular migration and invasion"

**Supplemental Figure 1. NDRG1 expression is stimulated by androgen treatment in LNCaP cells**

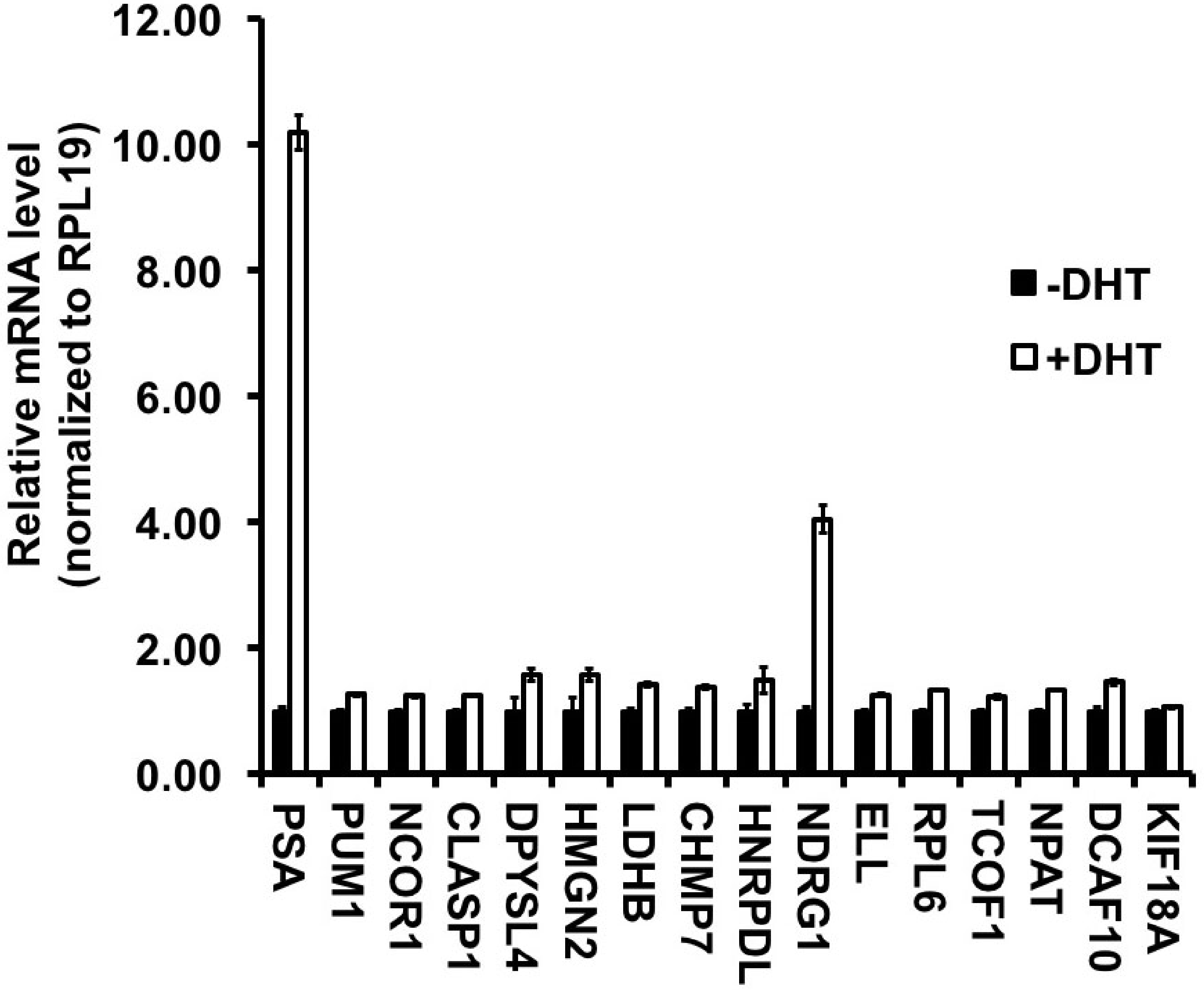

**Supplemental Figure 2. Supplemental Figures Associated with Figure 6**

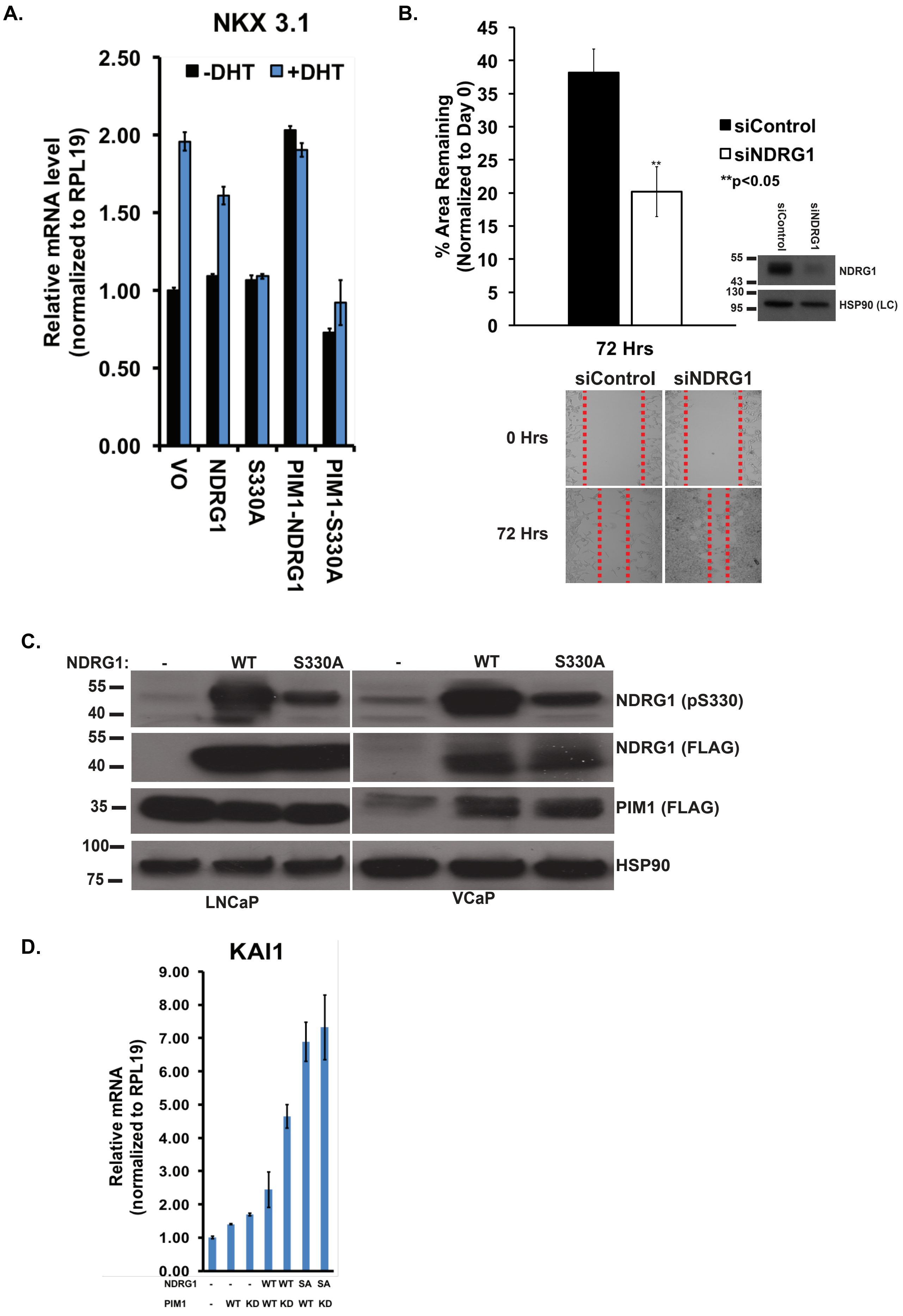

#### Supplemental Figure 3. Metascape Analysis of PIM1 Substrates

**A.**

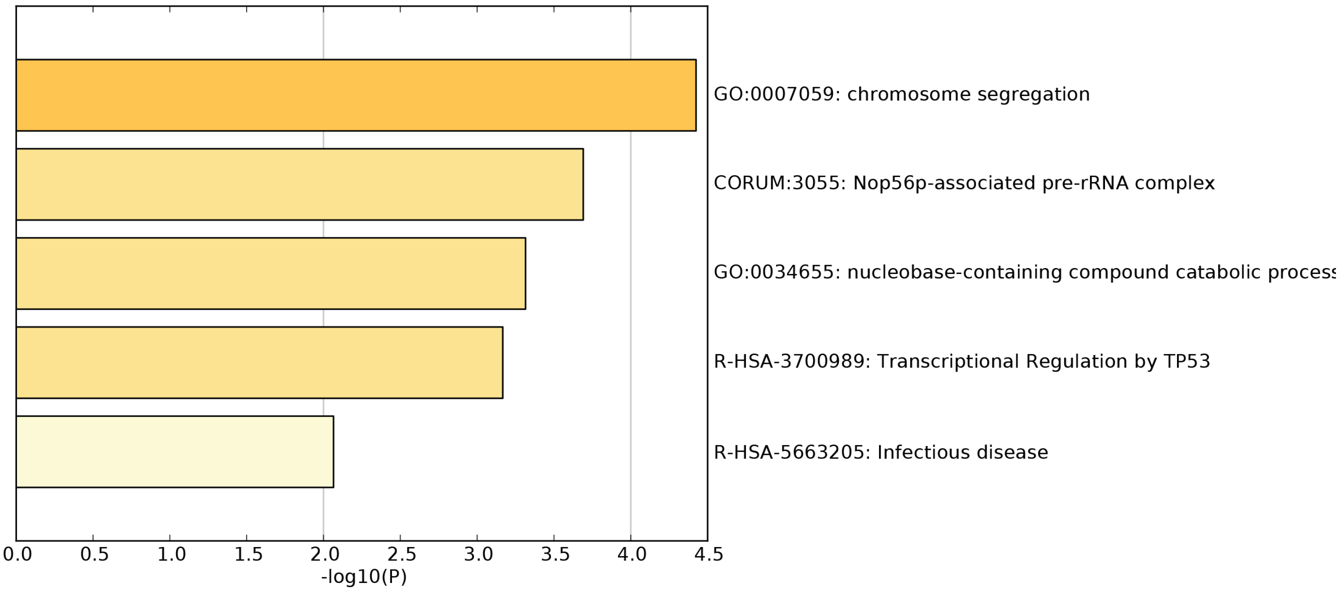

**B.**

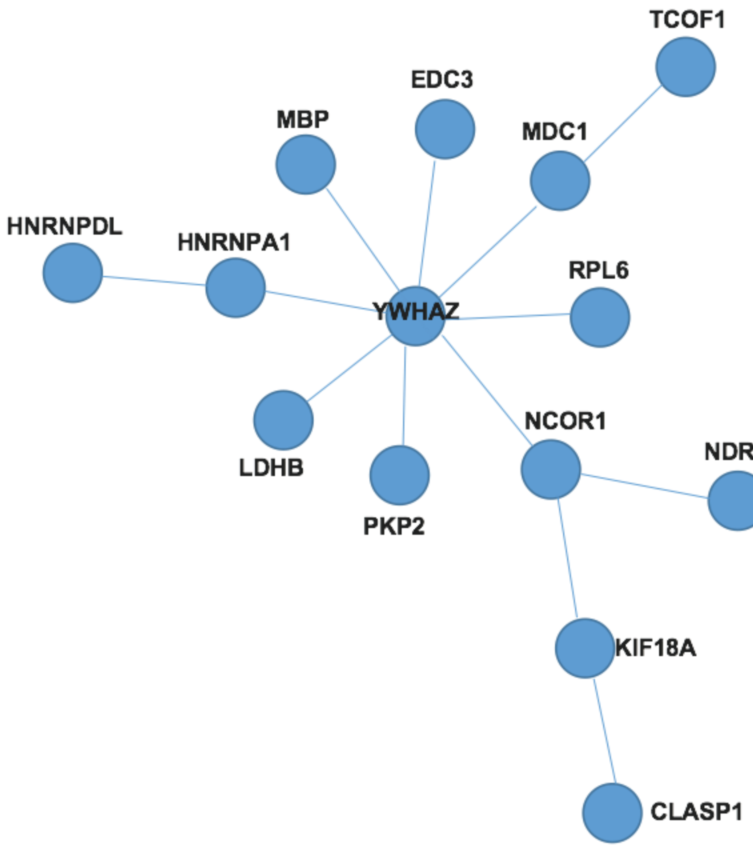

**Supplemental Table 1. PIM1 Gatekeeper Residue Alignment**

| **PIM1 Residue** | | | | | | | **120** | | | | | |
|---|---|---|---|---|---|---|---|---|---|---|---|---|
| ***H.s.* PIM1** | S | H | F | V | L | I | **L** | E | R | P | E | P |
| *H.s.* AMPK | T | D | F | F | M | V | **M** | E | Y | V | S | G |
| *H.s.* CDK2 | N | K | L | Y | L | V | **F** | E | F | L | H | Q |
| *M.m.* Jnk1 | Q | D | V | Y | I | V | **M** | E | L | M | D | A |
| *M.m.* Pka ca | S | N | L | Y | M | V | **M** | E | Y | V | P | G |
| *S.c.* CDC28 | H | K | L | Y | L | V | **F** | E | F | L | D | L |
| *S.c.* CLA4 | E | E | L | W | V | I | **M** | E | Y | M | E | G |

#### Supplemental Table 2. PIM1 Aggregate Substrate Identification

| UniPROT Accession ID | Protein | Identified Phosphosite | Peptides Identified (Exp.1) | Peptides Identified (Exp. 2) | Validated | Function |
|---|---|---|---|---|---|---|
| Q14671 | PUM1 | T112 | 32 | 27 | Yes | RNA-binding protein translation control |
| O75376 | NCOR1 | S1450 | 6 | 6 | No | transcriptional regulation/ co-repressor |
| Q7Z460 | CLASP1 | S647 | 6 | 2 | No | microtubule binding protein /stabilizes microtubules |
| O14531 | DPYSL4 | S537 | 4 | 1 | No | semaphorin signaling / cytoskeleton remodeling |
| P05204 | HMGN2 | S29 | 4 | 4 | No | alters interaction between DNA and histone octamer |
| P07195 | LDHB | T302 | 4 | 4 | No | synthesizes (S)-lactate from pyruvate |
| Q8WUX9 | CHPM7 | T408 | 4 | 8 | Yes | promotes nuclear envelope sealing/ spindle disassembly |
| O14979 | hnRNPDL | T62 | 2 | 4 | No | DNA/RNA binding protein: transcriptional repressor |
| Q92597 | NDRG1 | S330 | 2 | 9 | Yes | tumor suppressor/ cell trafficking |
| P55199 | ELL | T345 | 1 | 1 | No | RNA pol II elongation factor |
| Q02878 | RPL6 | T92 | 1 | 2 | No | component of the large 60s ribosomal subunit |
| Q13428 | TCOF1 | S1350 | 1 | 1 | No | regulator of RNA polymerase I |
| Q14207 | NPAT | T115 | 1 | 1 | No | transcriptional activator / cell cycle progression |
| Q5QP82 | DCAF10 | T348 | 1 | 1 | No | substrate receptor ubiquitin-protein ligase |
| Q8NI77 | KIF18A | T834 | 1 | 2 | Yes | microtubule-depolymerizer /chromosome congression |
| P63104 | YWHAZ | S64 | 4 | - | No | adapter protein/ regulates signaling |
| Q13442 | PDAP1 | S19 | 2 | - | No | enhances PDGFA-stimulated cell growth |
| Q14676 | MDC1 | S1113 | - | 2 | No | cell cycle arrest in response to DNA damage |
| Q7Z628 | NET1 | S21 | - | 2 | No | GEF for RhoA GTPase. |
| P59672 | ANKS1 | S887 | 1 | - | No | regulates EPHA8 receptor signaling to control cell migration |
| Q96F86 | EDC3 | S161 | 2 | - | No | mRNA degradation/ promotes mRNA de-capping |
| Q9H0H5 | RACGAP1 | T249 | - | 6 | No | myosin contractile ring formation during cytokinesis |
| P02686 | MBP | T229 | 1 | - | No | myelin membrane formation and stabilization |
| Q99959 | PKP2 | T320 | - | 1 | No | adherence junction maintenance |
| P09651 | hnRNPA1 | S199 | 7 | - | No | mRNA transport from nucleus to cytoplasm |

**Supplemental Table 3. PIM1 Substrate Phosphosite Alignment (All Substrates)**

| Protein | Phosphosite | -7 | -6 | -5 | -4 | -3 | -2 | -1 | 0 | 1 | 2 | 3 |
|---|---|---|---|---|---|---|---|---|---|---|---|---|
| PUM1 | T112 | N | S | K | H | R | W | P | S | G | D | N |
| NCOR1 | S1450 | T | V | R | S | R | H | T | S | V | V | S |
| CLASP1 | S647 | I | R | T | R | R | Q | S | S | G | S | A |
| DPYSL4 | S537 | P | V | R | N | L | H | Q | S | G | F | S |
| HMGN2 | S29 | Q | R | R | S | A | R | L | S | A | K | P |
| LDHB | T302 | I | L | N | A | R | G | L | T | S | V | I |
| CHMP7 | T408 | N | P | R | N | R | H | F | T | N | S | V |
| hnRNPDL | T62 | R | R | A | Q | R | H | V | T | A | Q | Q |
| NDRG1 | S330 | L | M | R | S | R | T | A | S | G | S | S |
| ELL | T345 | K | P | R | I | S | H | F | T | Q | R | A |
| RPL6 | T92 | K | K | E | K | V | L | A | T | V | T | K |
| TCOF1 | S1350 | E | S | R | K | R | K | L | S | G | D | Q |
| NPAT | T115 | S | Q | R | A | R | T | R | T | G | I | A |
| DCAF10 | T348 | I | L | R | A | R | R | T | T | S | S | S |
| KIF18A | T834 | A | K | R | K | R | K | L | T | S | S | T |
| YWHAZ* | S64 | S | S | W | R | V | V | S | S | I | E | Q |
| PDAP1* | S19 | G | R | A | R | Q | Y | T | S | P | E | E |
| MDC1* | S1113 | K | I | R | T | R | K | S | S | R | M | T |
| NET1* | S21 | R | R | R | S | R | R | A | S | G | L | S |
| ANKS1* | S887 | T | G | R | R | R | H | D | S | L | H | D |
| EDC3* | S161 | S | F | R | R | R | H | N | S | W | S | S |
| RACGAP1* | T249 | W | T | R | S | R | R | K | T | G | T | L |
| MBP* | T229 | K | N | I | V | T | P | R | T | P | P | S |
| PKP2* | T320 | S | G | R | R | A | H | L | T | V | G | Q |
| hnRNPA1* | S199 | S | Q | R | G | R | S | G | S | G | N | F |

Highlighted residues represent optimal, secondary, or tertiary residues based on **Table 2.** Red, basic residues. Green, hydrophobic residues. Blue, neutral polar residues. * **means substrate only identified in one replicate.**

TABLE 4A  Statistical Analyses of Prostate TMAs

**Description**

| | | N | Mean | Std. Deviation | Std. Error | 95% Confidence Interval | |
|---|---|---|---|---|---|---|---|
| | | | | | | Lower Bound | Upper Bound |
| NDRG1 PA330 | Control | 6 | 0.33 | 0.816 | 0.333 | -0.52 | 1.19 |
| | Gleason 3 | 19 | 0.42 | 1.017 | 0.233 | -0.07 | 0.91 |
| | Gleason 4 | 20 | 0.9 | 1.071 | 0.24 | 0.4 | 1.4 |
| | Gleason 5 | 29 | 1.14 | 1.217 | 0.226 | 0.68 | 1.6 |
| NDRG1 | Control | 6 | 2 | 0.894 | 0.365 | 1.06 | 2.94 |
| | Gleason 3 | 19 | 1.63 | 1.065 | 0.244 | 1.12 | 2.14 |
| | Gleason 4 | 20 | 1.7 | 1.129 | 0.252 | 1.17 | 2.23 |
| | Gleason 5 | 29 | 1.86 | 1.125 | 0.209 | 1.43 | 2.29 |
| | Total | 74 | 1.77 | 1.08 | 0.126 | 1.52 | 2.02 |

ANOVA

| | | Sum of Squares | df | Mean Square | F | Sig. |
|---|---|---|---|---|---|---|
| NDRG1 pS330 | Between Groups | 7.503 | 3 | 2.501 | 2.055 | 0.114 |
| | Within Groups | 85.213 | 70 | 1.217 | | |
| | Total | 92.716 | 73 | | | |
| NDRG1 | Between Groups | 1.025 | 3 | 0.342 | 0.285 | 0.836 |
| | Within Groups | 84.069 | 70 | 1.201 | | |
| | Total | 85.095 | 73 | | | |

**Correlations**

| | | Gleson_score | NDRG1 pS330 | NDRG1 |
|---|---|---|---|---|
| Gleson_score | Pearson Correlation | 1 | .249* | -0.002 |
| | Sig. (2-tailed) | | 0.032 | 0.987 |
| | N | 74 | 74 | 74 |
| NDRG1 pS330 | Pearson Correlation | .249* | 1 | .372** |
| | Sig. (2-tailed) | 0.032 | | 0.001 |
| | N | 74 | 74 | 74 |
| Total_NDRG1 | Pearson Correlation | -0.002 | .372** | 1 |
| | Sig. (2-tailed) | 0.987 | 0.001 | |
| | N | 74 | 74 | 74 |

* Correlation is significant at the 0.05 level (2-tailed).
** Correlation is significant at the 0.01 level (2-tailed).

Correlations

| | | | Gleson_score | NDRG1 pS330 | NDRG1 | PIM1 | AR |
|---|---|---|---|---|---|---|---|
| Spearman's rho | Gleson_score | Correlation Coefficie | 1 | .338** | 0.195 | -0.077 | -0.176 |
| | | Sig. (2-tailed) | . | 0.003 | 0.091 | 0.535 | 0.15 |
| | | N | 76 | 76 | 76 | 67 | 68 |

| | | | | | | |
|---|---|---|---|---|---|---|
| NDRG1 pS330 | Correlation Coefficie | .338** | 1 | .381** | 0.139 | 0.147 |
| | Sig. (2-tailed) | 0.003 | . | 0 | 0.261 | 0.233 |
| | N | 76 | 80 | 80 | 67 | 68 |
| NDRG1 | Correlation Coefficie | 0.195 | .381** | 1 | .303* | 0.23 |
| | Sig. (2-tailed) | 0.091 | 0 | . | 0.013 | 0.06 |
| | N | 76 | 80 | 80 | 67 | 68 |
| PIM1 | Correlation Coefficie | -0.077 | 0.139 | .303* | 1 | .376** |
| | Sig. (2-tailed) | 0.535 | 0.261 | 0.013 | . | 0.002 |
| | N | 67 | 67 | 67 | 67 | 67 |
| AR | Correlation Coefficie | -0.176 | 0.147 | 0.23 | .376** | 1 |
| | Sig. (2-tailed) | 0.15 | 0.233 | 0.06 | 0.002 | . |
| | N | 68 | 68 | 68 | 67 | 68 |

** Correlation is significant at the 0.01 level (2-tailed).

* Correlation is significant at the 0.05 level (2-tailed).

| Spearman r | AR | AR pS213 | PIM1 | NDRG1 | NDRG1 pS330 |
|---|---|---|---|---|---|
| AR441 | | 0.302038259 | 0.29732471 | 0.460469027 | 0.097310369 |
| P-AR S213 | 0.302038259 | | 0.602405453 | 0.313578966 | 0.366370197 |
| PIM1 | 0.29732471 | 0.602405453 | | 0.275309294 | 0.50376836 |
| NDRG11 | 0.460469027 | 0.313578966 | 0.275309294 | | 0.282972212 |
| p-NDRG1 | 0.097310369 | 0.366370197 | 0.50376836 | 0.282972212 | |

| p value | AR | P-AR S213 | PIM1 | NDRG1 | NDRG1 pS330 |
|---|---|---|---|---|---|
| AR441 | | 0.036939982 | 0.040140969 | 0.000990333 | 0.510551645 |
| P-AR S213 | 0.036939982 | | 5.89547E-06 | 0.029981134 | 0.010433755 |
| PIM1 | 0.040140969 | 5.89547E-06 | | 0.058238263 | 0.000262036 |
| NDRG11 | 0.000990333 | 0.029981134 | 0.058238263 | | 0.051313641 |
| p-NDRG1 | 0.510551645 | 0.010433755 | 0.000262036 | 0.051313641 | |

#### Table 4B Characteristics and Scoring of TMA for NDRG1 pS330, NDRG1 , PIM1 and AR

| catalog # | position | sex | age | organ | pathology | grade | stage | gleason_grade | gleason_score | tnm | type | IHC Score | | | |
|---|---|---|---|---|---|---|---|---|---|---|---|---|---|---|---|
| | | | | | | | | | | | | pNDRG1 | NDRG1 | PIM1 | AR |
| PR803a | A1 | M | 60 | Prostate | Adenocarcinoma | 1 | IV | 2 | 2+2 | T4N1M1 | Malignant | 0 | 1 | 1 | 2 |
| PR803a | A2 | M | 75 | Prostate | Adenocarcinoma | 1 | IV | 2 | 2+2 | T2N0M1c | Malignant | 0 | 1 | 3 | 1 |
| PR803a | A3 | M | 60 | Prostate | Adenocarcinoma | 3 | III | 4 | 5+4 | T3aN0M0 | Malignant | 0 | 1 | 3 | 2 |
| PR803a | A4 | M | 71 | Prostate | Adenocarcinoma | 2 | II | 3 | 3+3 | T2N0M0 | Malignant | 0 | 3 | 3 | 2 |
| PR803a | A5 | M | 66 | Prostate | Adenocarcinoma | 2 | IV | 3 | 3+3 | T3N1M1 | Malignant | 0 | 2 | 1 | 2 |
| PR803a | A6 | M | 67 | Prostate | Adenocarcinoma | 2 | IV | 3 | 2+3 | T2N0M1 | Malignant | 3 | 2 | 2 | 1 |
| PR803a | A7 | M | 72 | Prostate | Adenocarcinoma | 2 | II | 3 | 2+3 | T2N0M0 | Malignant | 0 | 0 | 2 | 2 |
| PR803a | A8 | M | 76 | Prostate | Adenocarcinoma | 2 | II | 3 | 2+3 | T2N0M0 | Malignant | 3 | 3 | 2 | 3 |
| PR803a | A9 | M | 71 | Prostate | Adenocarcinoma | 2 | II | 3 | 4+3 | T2N0M0 | Malignant | 0 | 1 | 1 | 3 |
| PR803a | A10 | M | 65 | Prostate | Adenocarcinoma | 2 | II | 3 | 4+3 | T2N0M0 | Malignant | 0 | 1 | 2 | 1 |
| PR803a | B1 | M | 64 | Prostate | Adenocarcinoma | 2 | I | 3 | 3+3 | T1N0M0 | Malignant | 0 | 2 | 2 | 1 |
| PR803a | B2 | M | 71 | Prostate | Adenocarcinoma | 3 | II | 4 | 5+4 | T2N0M0 | Malignant | 0 | 2 | 2 | 2 |
| PR803a | B3 | M | 71 | Prostate | Adenocarcinoma | 2 | I | 3 | 3+3 | T1N0M0 | Malignant | 0 | 1 | 2 | 1 |
| PR803a | B4 | M | 72 | Prostate | Adenocarcinoma (smooth muscle) | - | II | - | - | T2N0M0 | Malignant | 1 | 0 | 3 | 1 |
| PR803a | B5 | M | 70 | Prostate | Adenocarcinoma | 3 | II | 5 | 5+5 | T2N0M0 | Malignant | 1 | 3 | 3 | 2 |
| PR803a | B6 | M | 73 | Prostate | Adenocarcinoma | 2 | II | 3 | 4+3 | T2N0M0 | Malignant | 0 | 1 | 2 | 1 |
| PR803a | B7 | M | 70 | Prostate | Adenocarcinoma | 2 | IV | 3 | 4+3 | T2N1M1c | Malignant | 0 | 1 | 2 | 2 |
| PR803a | B8 | M | 65 | Prostate | Adenocarcinoma | 2 | II | 3 | 4+3 | T2N0M0 | Malignant | 2 | 3 | 3 | 2 |
| PR803a | B9 | M | 75 | Prostate | Adenocarcinoma | 2 | IV | 3 | 4+3 | T4N1M1 | Malignant | 2 | 3 | 2 | 2 |
| PR803a | B10 | M | 76 | Prostate | Adenocarcinoma (prostate tissue) | - | II | - | - | T2aN0M0 | Malignant | 0 | 0 | | 1 |
| PR803a | C1 | M | 64 | Prostate | Adenocarcinoma | 2 | IV | 3 | 4+3 | T3N0M1 | Malignant | 0 | 3 | 2 | 1 |
| PR803a | C2 | M | 74 | Prostate | Adenocarcinoma | 2--3 | IV | 4 | 5+3 | T4N1M1c | Malignant | 1 | 2 | 1 | 1 |
| PR803a | C3 | M | 62 | Prostate | Adenocarcinoma | 2 | IV | 3 | 4+3 | T3N1M1b | Malignant | 0 | 1 | 2 | 2 |
| PR803a | C4 | M | 64 | Prostate | Adenocarcinoma | 2--3 | II | 4 | 5+3 | T2aN0M0 | Malignant | 0 | 2 | 2 | 1 |
| PR803a | C5 | M | 58 | Prostate | Adenocarcinoma | 2 | II | 4 | 4+3 | T2N0M0 | Malignant | 0 | 3 | 3 | 2 |
| PR803a | C6 | M | 56 | Prostate | Adenocarcinoma | 2 | II | 3 | 4+3 | T2N0M0 | Malignant | 0 | 0 | 1 | 1 |
| PR803a | C7 | M | 62 | Prostate | Adenocarcinoma | 2 | II | 3 | 3+3 | T2N0M0 | Malignant | 1 | 3 | 2 | 1 |
| PR803a | C8 | M | 73 | Prostate | Adenocarcinoma | 2 | II | 3 | 3+3 | T2N0M0 | Malignant | 0 | 2 | 2 | 2 |
| PR803a | C9 | M | 71 | Prostate | Adenocarcinoma | 2 | II | 4 | 4+3 | T2N0M0 | Malignant | 0 | 0 | 1 | 1 |
| PR803a | C10 | M | 64 | Prostate | Adenocarcinoma | 2--3 | II | 4 | 3+5 | T2N0M0 | Malignant | 3 | 3 | 3 | 1 |
| PR803a | D1 | M | 70 | Prostate | Adenocarcinoma (sparse) | - | III | - | - | T3N0M0 | Malignant | 0 | 0 | | |
| PR803a | D2 | M | 73 | Prostate | Adenocarcinoma (prostate tissue) | - | II | - | - | T2N0M0 | Malignant | 0 | 0 | | |
| PR803a | D3 | M | 70 | Prostate | Adenocarcinoma | 2 | IV | 3 | 4+3 | T3N1M0 | Malignant | 0 | 3 | 3 | 3 |
| PR803a | D4 | M | 72 | Prostate | Adenocarcinoma | 3 | IV | 4 | 5+4 | T4N0M0 | Malignant | 1 | 1 | 1 | 1 |
| PR803a | D5 | M | 64 | Prostate | Adenocarcinoma | 3 | IV | 4 | 5+4 | T3N0M1a | Malignant | 3 | 2 | 3 | 3 |
| PR803a | D6 | M | 80 | Prostate | Adenocarcinoma | 2 | IV | 3 | 4+3 | T4N1M1c | Malignant | 0 | 2 | 1 | 1 |
| PR803a | D7 | M | 69 | Prostate | Adenocarcinoma | 2 | III | 3 | 4+3 | T3bN0M0 | Malignant | 0 | 3 | 2 | 1 |
| PR803a | D8 | M | 73 | Prostate | Adenocarcinoma | 3 | III | 4 | 5+4 | T3N1M1 | Malignant | 3 | 2 | 3 | 2 |
| PR803a | D9 | M | 73 | Prostate | Adenocarcinoma | 2--3 | IV | 4 | 5+3 | T3N1M1b | Malignant | 0 | 1 | 1 | 1 |
| PR803a | D10 | M | 65 | Prostate | Adenocarcinoma | 2--3 | IV | 4 | 3+5 | T2N1M1 | Malignant | 2 | 2 | 3 | 2 |
| PR803a | E1 | M | 82 | Prostate | Adenocarcinoma | 2 | IV | 3 | 4+3 | T3N2M1 | Malignant | 2 | 1 | 1 | 1 |
| PR803a | E2 | M | 51 | Prostate | Adenocarcinoma | 2 | II | 3 | 4+3 | T2N0M0 | Malignant | 3 | 3 | 1 | 1 |
| PR803a | E3 | M | 72 | Prostate | Adenocarcinoma | 2--3 | III | 4 | 5+3 | T3N0M0 | Malignant | 1 | 0 | 3 | 2 |

| | | | | | | | | | | | | | | | |
|---|---|---|---|---|---|---|---|---|---|---|---|---|---|---|---|
| PR803a | E4 | M | 66 | Prostate | Adenocarcinoma | 2 | III | 3 | 4+3 | T3aN0M0 | Malignant | 3 | 3 | 1 | 2 |
| PR803a | E5 | M | 20 | Prostate | Adenocarcinoma | 2 | III | 3 | 3+3 | T3N0M0 | Malignant | 0 | 1 | 2 | 1 |
| PR803a | E6 | M | 61 | Prostate | Adenocarcinoma | 3 | IV | 4 | 5+4 | T3N1M0 | Malignant | 0 | 3 | 1 | 1 |
| PR803a | E7 | M | 64 | Prostate | Adenocarcinoma | 2 | II | 3 | 4+3 | T2N0M0 | Malignant | 2 | 2 | 3 | 3 |
| PR803a | E8 | M | 60 | Prostate | Adenocarcinoma | 2 | IV | 3 | 4+3 | T3N1M0 | Malignant | 2 | 3 | 3 | 3 |
| PR803a | E9 | M | 68 | Prostate | Adenocarcinoma | 2 | II | 3 | 4+3 | T2N0M0 | Malignant | 0 | 0 | 1 | 2 |
| PR803a | E10 | M | 82 | Prostate | Adenocarcinoma | 3 | II | 5 | 5+5 | T2N0M0 | Malignant | 3 | 3 | 3 | 3 |
| PR803a | F1 | M | 78 | Prostate | Adenocarcinoma | 3 | IV | 4 | 5+4 | T3N2M1 | Malignant | 2 | 2 | 1 | 1 |
| PR803a | F2 | M | 87 | Prostate | Adenocarcinoma | 3 | II | 5 | 5+5 | T2N0M0 | Malignant | 0 | 1 | 1 | 1 |
| PR803a | F3 | M | 81 | Prostate | Adenocarcinoma | 3 | III | 4 | 5+4 | T3aN0M0 | Malignant | 0 | 2 | 3 | 1 |
| PR803a | F4 | M | 76 | Prostate | Adenocarcinoma | 3 | IV | 5 | 5+5 | T3N1M1b | Malignant | 0 | 0 | 1 | 1 |
| PR803a | F5 | M | 73 | Prostate | Adenocarcinoma | 3 | IV | 4 | 5+4 | T4N1M1c | Malignant | 2 | 3 | 2 | 1 |
| PR803a | F6 | M | 78 | Prostate | Adenocarcinoma | 3 | IV | 5 | 5+5 | T4N1M1 | Malignant | 3 | 2 | 1 | 1 |
| PR803a | F7 | M | 40 | Prostate | Adenocarcinoma | 3 | IV | 4 | 5+4 | T2N1M1b | Malignant | 0 | 0 | 1 | 2 |
| PR803a | F8 | M | 69 | Prostate | Adenocarcinoma | 2 | II | 3 | 4+3 | T2N0M0 | Malignant | 0 | 0 | 1 | 1 |
| PR803a | F9 | M | 62 | Prostate | Adenocarcinoma | 3 | II | 5 | 5+5 | T2N0M0 | Malignant | 0 | 3 | 2 | 1 |
| PR803a | F10 | M | 67 | Prostate | Adenocarcinoma | 3 | II | 5 | 5+5 | T2N0M0 | Malignant | 1 | 3 | 2 | 1 |
| PR803a | G1 | M | 64 | Prostate | Adenocarcinoma | 3 | II | 5 | 5+5 | T2N0M0 | Malignant | 0 | 3 | 3 | 2 |
| PR803a | G2 | M | 77 | Prostate | Adenocarcinoma | 2 | II | 3 | 4+3 | T2N0M0 | Malignant | 2 | 1 | 1 | 1 |
| PR803a | G3 | M | 26 | Prostate | Adenocarcinoma (prostate tissue) | - | II | - | - | T2N0M0 | Malignant | 0 | 1 | 2 | 1 |
| PR803a | G4 | M | 69 | Prostate | Adenocarcinoma | 3 | IV | 5 | 5+5 | T4N0M0 | Malignant | 1 | 1 | 2 | 1 |
| PR803a | G5 | M | 73 | Prostate | Adenocarcinoma | 2 | III | 3 | 4+3 | T3N0M0 | Malignant | 0 | 2 | 2 | 1 |
| PR803a | G6 | M | 82 | Prostate | Adenocarcinoma | 2--3 | II | 4 | 5+3 | T2N0M0 | Malignant | 0 | 2 | 1 | 1 |
| PR803a | G7 | M | 63 | Prostate | Adenocarcinoma | 2--3 | IV | 4 | 5+3 | T2N1M1b | Malignant | 1 | 1 | 1 | 1 |
| PR803a | G8 | M | 55 | Prostate | Adenocarcinoma | 3 | II | 4 | 5+4 | T2N0M0 | Malignant | 1 | 1 | 1 | 1 |
| PR803a | G9 | M | 75 | Prostate | Adenocarcinoma with necrosis | 3 | III | 4 | 5+4 | T3N0M0 | Malignant | 1 | 1 | 3 | 1 |
| PR803a | G10 | M | 72 | Prostate | Adenocarcinoma | 3 | IV | 4 | 5+4 | T2N0M1 | Malignant | 0 | 0 | 3 | 1 |
| PR803a | H1 | M | 60 | Prostate | Adenocarcinoma (sparse) | - | II | - | - | T2N0M0 | Malignant | 2 | 2 | | |
| PR803a | H2 | M | 72 | Prostate | Leiomyosarcoma | - | II | - | - | T2N0M0 G | Malignant | 0 | 0 | | |
| PR803a | H3 | M | 67 | Prostate | Pleomorphic leiomyosarcoma | - | IIIB | - | - | T2N0M0 G | Malignant | 0 | 0 | | |
| PR803a | H4 | M | 27 | Prostate | Hyperplasia | - | - | - | - | - | Hyperplasi | 0 | 2 | | |
| PR803a | H5 | M | 35 | Prostate | Cancer adjacent normal prostate tissue | - | - | - | - | - | NAT | 0 | 2 | | |
| PR803a | H6 | M | 21 | Prostate | Cancer adjacent normal prostate tissue | - | - | - | - | - | NAT | 2 | 3 | | |
| PR803a | H7 | M | 45 | Prostate | Normal prostate tissue | - | - | - | - | - | Normal | 0 | 3 | | |
| PR803a | H8 | M | 37 | Prostate | Normal prostate tissue | - | - | - | - | - | Normal | 0 | 2 | | |
| PR803a | H9 | M | 33 | Prostate | Normal prostate tissue | - | - | - | - | - | Normal | 0 | 1 | | |
| PR803a | H10 | M | 43 | Prostate | Normal prostate tissue | - | - | - | - | - | Normal | 0 | 1 | | |

### Table 4C Characteristics and Scoring of TMA for AR, AR pS213 and PIM1

| PR807a | catalognur | position | sex | age | organ | pathology | grade | stage | gleason_gr | gleason_sc | tnm | type | IHC Score | | |
|---|---|---|---|---|---|---|---|---|---|---|---|---|---|---|---|
| | | | | | | | | | | | | | AR | ARpS213 | PIM1 |
| A1 | PR807b | A1 | M | 43 | Prostate | Prostatic tissue | - | - | - | - | - | | 1 | 1 | 0 |
| A4 | PR807b | A2 | M | 35 | Prostate | Prostatic tissue | - | - | - | - | - | | 1 | 1 | 1 |
| A6 | PR807b | A3 | M | 28 | Prostate | Prostatic tissue | - | - | - | - | - | | 1 | 0 | 0 |
| B1 | PR807b | A4 | M | 27 | Prostate | Adjacent normal prostatic tissue | - | - | - | - | - | | 1 | 1 | 1 |
| B2 | PR807b | A5 | M | 25 | Prostate | Adjacent normal prostatic tissue | - | - | - | - | - | | 1 | 1 | 0 |
| B3 | PR807b | A6 | M | 21 | Prostate | Adjacent normal prostatic tissue | - | - | - | - | - | | 1 | 1 | 0 |
| B4 | PR807b | A7 | M | 64 | Prostate | Adjacent normal prostatic tissue | - | - | - | - | - | | 1 | 2 | 0 |
| B5 | PR807b | A8 | M | 84 | Prostate | Adjacent normal prostatic tissue | - | - | - | - | - | | 1 | 2 | 0 |
| B6 | PR807b | A9 | M | 80 | Prostate | Adjacent normal prostatic tissue | - | - | - | - | - | | 1 | 1 | 0 |
| B7 | PR807b | A10 | M | 77 | Prostate | Adjacent normal prostatic tissue | - | - | - | - | - | | 1 | 2 | 0 |
| B8 | PR807b | B1 | M | 71 | Prostate | Adenocarcinoma | 1 | II | 2 | 2+2 | T2N0M0 | Tumor | 2 | 3 | 3 |
| B9 | PR807b | B2 | M | 64 | Prostate | Adenocarcinoma | 1 | I | 2 | 2+3 | T1N0M0 | Tumor | 1 | 2 | 1 |
| B10 | PR807b | B3 | M | 66 | Prostate | Adenocarcinoma | 1 | IV | 2 | 2+2 | T3N1M1 | Tumor | 1 | 1 | 0 |
| C1 | PR807b | B4 | M | 76 | Prostate | Adenocarcinoma | 1 | II | 2 | 2+3 | T2aN0M0 | Tumor | 2 | 3 | 2 |
| C2 | PR807b | B5 | M | 72 | Prostate | Adenocarcinoma | 1 | II | 2 | 2+3 | T2N0M0 | Tumor | 1 | 1 | 0 |
| C3 | PR807b | B6 | M | 69 | Prostate | Adenocarcinoma | 1--2 | III | 2 | 2+3 | T3N0M0 | Tumor | 1 | 1 | 1 |
| C4 | PR807b | B7 | M | 71 | Prostate | Adenocarcinoma | 1 | II | 2 | 2+2 | T2N0M0 | Tumor | 2 | 3 | 1 |
| C5 | PR807b | B8 | M | 65 | Prostate | Adenocarcinoma | 1 | II | 2 | 2+3 | T2N0M0 | Tumor | 2 | 2 | 1 |
| C6 | PR807b | B9 | M | 65 | Prostate | Adenocarcinoma | 2 | II | 3 | 3+4 | T2N0M0 | Tumor | 1 | 1 | 0 |
| C7 | PR807b | B10 | M | 64 | Prostate | Adenocarcinoma | 2 | IV | 3 | 3+3 | T3N0M1b | Tumor | 2 | 1 | 0 |
| C8 | PR807b | C1 | M | 75 | Prostate | Adenocarcinoma | 2 | IV | 3 | 3+3 | T4N1M1 | Tumor | 2 | 1 | 3 |
| C9 | PR807b | C2 | M | 70 | Prostate | Adenocarcinoma | 2 | II | 3 | 3+4 | T2N0M0 | Tumor | 2 | 2 | 3 |
| C10 | PR807b | C3 | M | 58 | Prostate | Adenocarcinoma | 2 | II | 3 | 3+4 | T2N0M0 | Tumor | 2 | 1 | 3 |
| D1 | PR807b | C4 | M | 61 | Prostate | Adenocarcinoma | 2 | III | 3 | 3+3 | T3N1M0 | Tumor | 2 | 1 | 3 |
| D2 | PR807b | C5 | M | 70 | Prostate | Adenocarcinoma | 2 | III | 3 | 3+4 | T3N0M0 | Tumor | 2 | 1 | 2 |
| D3 | PR807b | C6 | M | 20 | Prostate | Adenocarcinoma | 3 | III | 4 | 4+5 | T3N0M0 | Tumor | 1 | 1 | 1 |
| D4 | PR807b | C7 | M | 62 | Prostate | Adenocarcinoma | 2 | IV | 3 | 3+3 | T3N1M1b | Tumor | 1 | 1 | 0 |
| D5 | PR807b | C8 | M | 64 | Prostate | Adenocarcinoma | 2 | II | 3 | 3+4 | T2aN0M0 | Tumor | 1 | 1 | 2 |
| D6 | PR807b | C9 | M | 73 | Prostate | Adenocarcinoma | 2 | II | 3 | 3+3 | T2N0M0 | Tumor | 1 | 1 | 0 |
| D7 | PR807b | C10 | M | 64 | Prostate | Adenocarcinoma | 2 | IV | 3 | 3+3 | T3N0M1 | Tumor | 1 | 3 | 3 |
| D8 | PR807b | D1 | M | 64 | Prostate | Adenocarcinoma | 2 | II | 3 | 3+4 | T2N0M0 | Tumor | 1 | 2 | 2 |
| D9 | PR807b | D2 | M | 82 | Prostate | Adenocarcinoma | 2 | II | 3 | 3+3 | T2N0M0 | Tumor | 3 | 3 | 3 |
| E1 | PR807b | D3 | M | 70 | Prostate | Adenocarcinoma | 2 | IV | 3 | 3+3 | T2N1M1c | Tumor | 2 | 0 | 0 |
| E2 | PR807b | D4 | M | 60 | Prostate | Adenocarcinoma | 2 | IV | 3 | 3+3 | T3N1M0 | Tumor | 3 | 2 | 3 |
| E3 | PR807b | D5 | M | 64 | Prostate | Adenocarcinoma | 2 | II | 3 | 3+4 | T2N0M0 | Tumor | 2 | 2 | 2 |
| E4 | PR807b | D6 | M | 72 | Prostate | Adenocarcinoma | 2 | III | 4 | 4+3 | T3N0M0 | Tumor | 2 | 1 | 0 |
| E5 | PR807b | D7 | M | 78 | Prostate | Adenocarcinoma | 2 | IV | 4 | 4+3 | T4N1M1b | Tumor | 2 | 0 | 1 |
| E6 | PR807b | D8 | M | 76 | Prostate | Adenocarcinoma | 2 | III | 4 | 4+4 | T3N1M0 | Tumor | 1 | 0 | 0 |
| E7 | PR807b | D9 | M | 72 | Prostate | Adenocarcinoma | 2 | III | 4 | 4+4 | T3N0M0 | Tumor | 2 | 0 | 0 |
| E8 | PR807b | D10 | M | 70 | Prostate | Adenocarcinoma | 3 | II | 4 | 4+5 | T2bN0M0 | Tumor | 3 | 3 | 3 |
| E9 | PR807b | E1 | M | 65 | Prostate | Adenocarcinoma | 2 | IV | 3 | 4+2 | T2N1M1 | Tumor | 1 | 2 | 3 |
| E10 | PR807b | E2 | M | 69 | Prostate | Adenocarcinoma | 3 | III | 4 | 4+5 | T4N0M0 | Tumor | 3 | 3 | 3 |
| F1 | PR807b | E3 | M | 80 | Prostate | Adenocarcinoma | 2 | IV | 3 | 3+3 | T4N1M1c | Tumor | 1 | 1 | 1 |
| F3 | PR807b | E4 | M | 82 | Prostate | Adenocarcinoma | 2 | IV | 4 | 3+5 | T3N2M1 | Tumor | 1 | 2 | 2 |
| F4 | PR807b | E5 | M | 73 | Prostate | Adenocarcinoma | 3 | IV | 4 | 4+5 | T3N1M1b | Tumor | 2 | 2 | 1 |
| F5 | PR807b | E6 | M | 62 | Prostate | Adenocarcinoma | 3 | II | 5 | 5+5 | T2N0M0 | Tumor | 2 | 1 | 0 |
| F6 | PR807b | E7 | M | 60 | Prostate | Adenocarcinoma (sparse) | 3 | III | 5 | 5+5 | T3aN0M0 | Tumor | 1 | 1 | 1 |

| | | | | | | | | | | | | | | | | | |
|---|---|---|---|---|---|---|---|---|---|---|---|---|---|---|---|---|---|
| F7 | PR807b | E8 | M | 26 | Prostate | Adenocarcinoma | - | | II | - | | - | T2N0M0 | Tumor | 1 | 0 | 1 |
| F8 | PR807b | E9 | M | 87 | Prostate | Adenocarcinoma | | 3 | II | | 5 | 5+5 | T2N0M0 | Tumor | 1 | 0 | 0 |
| F9 | PR807b | E10 | M | 75 | Prostate | Adenocarcinoma | | 3 | IV | | 5 | 5+5 | T2N1M1c | Tumor | 2 | 0 | 0 |
| F10 | PR807b | F1 | M | 64 | Prostate | Adenocarcinoma | 2--3 | | II | | 4 | 4+5 | T2N0M0 | Tumor | 1 | 1 | 2 |
| | PR807b | F2 | M | 60 | Prostate | Adenocarcinoma | | 2 | IV | | 3 | 4+3 | T3N1M1b | Tumor | 1 | 0 | 1 |
| | PR807b | F3 | M | 73 | Prostate | Adenocarcinoma | | 3 | IV | | 5 | 5+5 | T3N1M1c | Tumor | 3 | 2 | 1 |
| | PR807b | F4 | M | 69 | Prostate | Adenocarcinoma | | 2 | II | | 3 | 4+3 | T2N0M0 | Tumor | 3 | 3 | 1 |
| | PR807b | F5 | M | 63 | Prostate | Adenocarcinoma | | 3 | IV | | 5 | 5+5 | T2N1M1b | Tumor | 3 | 2 | 3 |
| | PR807b | F6 | M | 55 | Prostate | Adenocarcinoma | | 3 | II | | 4 | 4+5 | T2N0M0 | Tumor | 2 | 0 | 0 |
| | PR807b | F7 | M | 82 | Prostate | Adenocarcinoma | | 3 | II | | 5 | 5+5 | T2N0M0 | Tumor | 2 | 0 | 1 |
| | PR807b | F8 | M | 73 | Prostate | Adenocarcinoma | | 3 | IV | | 5 | 5+5 | T4N1M1c | Tumor | 3 | 2 | 2 |
| | PR807b | F9 | M | 75 | Prostate | Adenocarcinoma with necrosis | | 3 | III | | 5 | 5+5 | T3N0M0 | Tumor | 3 | 1 | 2 |
| | PR807b | F10 | M | 60 | Prostate | Adenocarcinoma | | 3 | II | | 5 | 5+5 | T2N0M0 | Tumor | 2 | 1 | 0 |
| | PR807b | G1 | M | 62 | Prostate | Hyperplasia of prostate | - | | - | - | | - | - | | 1 | | |
| | PR807b | G2 | M | 78 | Prostate | Hyperplasia of prostate | - | | - | - | | - | - | | 1 | | |
| | PR807b | G3 | M | 73 | Prostate | Hyperplasia of prostate | - | | - | - | | - | - | | 1 | | |
| | PR807b | G4 | M | 70 | Prostate | Hyperplasia of prostate | - | | - | - | | - | - | | 1 | | |
| | PR807b | G5 | M | 70 | Prostate | Hyperplasia of prostate | - | | - | - | | - | - | | 1 | | |
| | PR807b | G6 | M | 73 | Prostate | Hyperplasia of prostate | - | | - | - | | - | - | | 1 | | |
| | PR807b | G7 | M | 74 | Prostate | Hyperplasia of prostate | - | | - | - | | - | - | | 1 | | |
| | PR807b | G8 | M | 81 | Prostate | Hyperplasia of prostate | - | | - | - | | - | - | | 1 | | |
| | PR807b | G9 | M | 72 | Prostate | Hyperplasia of prostate | - | | - | - | | - | - | | 1 | | |
| | PR807b | G10 | M | 78 | Prostate | Hyperplasia of prostate | - | | - | - | | - | - | | 1 | | |
| | PR807b | H1 | M | 66 | Prostate | Hyperplasia of prostate | - | | - | - | | - | - | | 1 | | |
| | PR807b | H2 | M | 79 | Prostate | Hyperplasia of prostate | - | | - | - | | - | - | | 1 | | |
| | PR807b | H3 | M | 65 | Prostate | Hyperplasia of prostate | - | | - | - | | - | - | | 1 | | |
| | PR807b | H4 | M | 81 | Prostate | Hyperplasia of prostate | - | | - | - | | - | - | | 1 | | |
| | PR807b | H5 | M | 66 | Prostate | Hyperplasia of prostate | - | | - | - | | - | - | | 1 | | |
| | PR807b | H6 | M | 67 | Prostate | Hyperplasia of prostate | - | | - | - | | - | - | | 1 | | |
| | PR807b | H7 | M | 72 | Prostate | Hyperplasia of prostate | - | | - | - | | - | - | | 1 | | |
| | PR807b | H8 | M | 56 | Prostate | Hyperplasia of prostate | - | | - | - | | - | - | | 1 | | |
| | PR807b | H9 | M | 76 | Prostate | Hyperplasia of prostate | - | | - | - | | - | - | | 1 | | |
| | PR807b | H10 | M | 68 | Prostate | Hyperplasia of prostate | - | | - | - | | - | - | | 1 | | |

**Supplemental Table 5 . Analysis of Phosphopeptides Corresponding to PIM1 Substrates from Drake et al. 2016 Phosphoproteomic Study**

| Phosphopeptide | Phosphosite | Gene Name | UniProt ID | Mean Difference (NonLog) Met vs. Primary |
|---|---|---|---|---|
| DpSLTGSSDLYK | pS709 | PUM1 | Q14671 | 0.92 |
| YETPSDAIEVIpSPApSSPAPPQEK | pS1977, pS1980 | NCOR1 | O75376 | 10.12 |
| pSPGSISYLPSFFTK | pS2184 | NCOR1 | O75376 | 3.30 |
| EpSPVSAPLEGLICR | pS1263 | NCOR1 | O75376 | 2.17 |
| AQLpSPGIYDDTSAR | pS1472 | NCOR1 | O75376 | 0.57 |
| ADpSVDVEVR | pS821 | NCOR1 | O75376 | 0.97 |
| NSDSIVSLPQpSDR | pS552 | CLASP1 | Q7Z460 | 2.98 |
| IpSDAELEAELEK | pS417 | CHMP7 | Q8WUX9 | 0.51 |
| VFVGGLpSPDTSEEQIK | pS241 | HNRPDL | O14979 | 2.05 |
| TApSGSSVTpSLDGTR | pS330, pS336 | NDRG1 | Q92597 | 7.73 |
| TASGpSpSVTSLDGTR | pS332, pS333 | NDRG1 | Q92597 | 6.67 |
| DNPpSPEPQLDDIK | pS165 | EAF1 | Q96JC9 | 7.70 |
| LDpSpSPSVSSTLAAK | pS1227, pS1228 | TCOF1 | Q13428 | 2.85 |
| LGAGEGGEApSVpSPEK | pS1376, pS1378 | TCOF1 | Q13428 | 6.32 |
| pSPAGPAATPAQAQAASTPR | pS967 | TCOF1 | Q13428 | 5.39 |
| SLEVGpSpYPILR | pS338, pY339 | DCAF10 | Q5QP82 | 0.17 |
| TAFDEAIAELDTLSEEpSYK | pS210 | YWHAZ | P63104 | 7.73 |
| DICNDVLpSLLEK | pS99 | YWHAZ | P63104 | 6.85 |
| TAFDEAIAELDTLpSEESYK | pS207 | YWHAZ | P63104 | 1.34 |
| SLDpSDEpSEDEEDDYQQK | pS60, pS63 | PDAP1 | Q13442 | 15.06 |
| MQSLpSLNK | pS178 | PDAP1 | Q13442 | 0.35 |
| MQpSLSLNK | pS176 | PDAP1 | Q13442 | 0.18 |
| DQPPFGDpSDDpSVEADK | pS495, pS498 | MDC1 | Q14676 | 7.03 |
| DpSDpTDVEEEELPVENR | pS453, pT455 | MDC1 | Q14676 | 3.26 |
| AQPFGFIDpSDpTDAEEER | pS329, pT331 | MDC1 | Q14676 | 2.98 |
| ESEDpSETQPFDTHLEAYGPCLpSPPR | pS763, pS780 | MDC1 | Q14676 | 1.59 |
| pSQLPAEGDAGAEWAAAVLK | pS598 | MDC1 | Q14676 | 1.28 |
| NQLVpTPEPTSR | pT1589 | MDC1 | Q14676 | 1.04 |
| pSQApSTTVDINTQVEK | pS513, pS516 | MDC1 | Q14676 | 0.67 |
| SQASTTVDINpTQVEK | pT523 | MDC1 | Q14676 | 0.43 |
| DQPPFGDpSDDpSVEADK | pS495, pS498 | MDC1 | Q14676 | 7.03 |
| SEpSLSNCSIGK | pS647 | ANKS1A | Q92625 | 1.45 |
| SQDVAVpSPQQQQCSK | pS131 | EDC3 | Q96F86 | 0.93 |
| pSIGSAVDQGNESIVAK | pS203 | RACGAP1 | Q9H0H5 | 2.47 |
| LEIpSPDpSSPER | pS151, pS154 | PKP2 | Q99959 | 1.57 |
| LEISPDpSpSPER | pS154, pS155 | PKP2 | Q99959 | 1.15 |
| pSEIVGVSR | pS197 | PKP2 | Q99959 | 0.34 |
| pSMGNLLEK | pS251 | PKP2 | Q99959 | 0.09 |
| pSKSEpSPKEPEQLR | pS2, pS6 | HNRNPA1 | P09651 | 4.86 |

#### Supplemental Figure 6. qPCR Primer List from Study

| | |
|---|---|
| **RPL6** | |
| Forward Primer | ATTCCCGATCTGCCATGTATTC |
| Reverse Primer | TACCGCCGTTCTTGTCACC |
| **ELL** | |
| Forward Primer | GATAGGAGCTACGGGCTGTC |
| Reverse Primer | TTCCTTGAAATCGGATAGATGGC |
| **HNRNPDL** | |
| Forward Primer | TCCGCTCCCGCTACTTTAG |
| Reverse Primer | CCTGCGCCCTCCCTTTATAG |
| **TCOF1** | |
| Forward Primer | CGGGAGCTACTTCCCCTGAT |
| Reverse Primer | CAGAAGGGTTACGGGCTGAG |
| **LDHB** | |
| Forward Primer | TGGTATGGCGTGTGCTATCAG |
| Reverse Primer | TTGGCGGTCACAGAATAATCTTT |
| **DCAF10** | |
| Forward Primer | GGACAATTTTCGCACCATGAC |
| Reverse Primer | ACAGCCGGTTATCAAGAAATCTG |
| **KIF10A** | |
| Forward Primer | TGCTGGGAAGACCCACACTAT |
| Reverse Primer | GCTGGTGTAAAGTAAGTCCATGA |
| **CHMP7** | |
| Forward Primer | AAGCCTCTCAAGTGGACTCTT |
| Reverse Primer | ACAGACGATACACCTCCTCAG |
| **NCOR1** | |
| Forward Primer | ACACCGCAGTATTGTCCAAAT |
| Reverse Primer | CACCTGGTTTGTCTTGATGTTCT |
| **PUM1** | |
| Forward Primer | ATGAGCGTTGCATGTGTCTTG |
| Reverse Primer | GTAGTCCACCATAGCGTCGTC |
| **CLASP1** | |
| Forward Primer | CTGTCTCTGCCTTATAGCAACAC |
| Reverse Primer | CATCTCGAACCTGGCTGTTTG |
| **DPYSL4** | |
| Forward Primer | CGTGAATGACGACCAGTCCTT |
| Reverse Primer | ACGATGAGGTTTTCTCCGATTTG |
| **NPAT** | |
| Forward Primer | AGAGCCCGAACGAGAACTG |
| Reverse Primer | GGTAAAGTGAGCAACTCTGCAC |
| **ELL** | |
| Forward Primer | GATAGGAGCTACGGGCTGTC |
| Reverse Primer | TTCCTTGAAATCGGATAGATGGC |
| **NDRG1** | |
| Forward Primer | CTCCTGCAAGAGTTTGATGTCC |
| Reverse Primer | TCATGCCGATGTCATGGTAGG |
| **PSA** | |
| Forward Primer | GTGTGTGGACCTCCATGTTATT |
| Reverse Primer | CCACTCACCTTTCCCCTCAAG |
| **Nkx3.1** | |
| Forward Primer | CCCACACTCAGGTGATCGAG |
| Reverse Primer | GAGCTGCTTTCGCTTAGTCTT |
